# Epstein-Barr virus reprograms human B-lymphocytes immediately in the pre-latent phase of infection

**DOI:** 10.1101/503268

**Authors:** Paulina Mrozek-Gorska, Alexander Buschle, Dagmar Pich, Thomas Schwarzmayr, Ron Fechtner, Antonio Scialdone, Wolfgang Hammerschmidt

**Affiliations:** Research Unit Gene Vectors, Helmholtz Zentrum München, German Research Center for Environmental Health and German Center for Infection Research (DZIF), Partner site Munich Marchioninistr. 25 D-81377 Munich, Germany; Institute of Human Genetics, Helmholtz Zentrum München, German Research Center for Environmental Health Ingolstaedter Landstr. 1 D-85764 Neuherberg, Germany; Institute of Computational Biology, Helmholtz Zentrum München, German Research Center for Environmental Health Ingolstaedter Landstr. 1 D-85764 Neuherberg, Germany; Institute of Epigenetics and Stem Cells, Helmholtz Zentrum München, German Research Center for Environmental Health Marchioninistr. 25 D-81377 Munich, Germany; Institute of Functional Epigenetics, Helmholtz Zentrum München, German Research Center for Environmental Health Ingolstaedter Landstr. 1 D-85764 Neuherberg, Germany

## Abstract

Epstein-Barr virus (EBV) is a human tumor virus and a model of herpesviral latency. The virus efficiently infects resting human B-lymphocytes and induces their continuous proliferation in vitro, which mimics certain aspects of EBV’s oncogenic potential in vivo. This seminal finding was made 50 years ago, but how EBV activates primary human B-lymphocytes and how lymphoblastoid cell lines (LCLs) evolve from the EBV-infected lymphocytes is uncertain. We conducted a systematic time-resolved longitudinal study of cellular functions and transcriptional profiles of newly infected naïve primary B-lymphocytes. EBV reprograms these human cells comprehensively and globally. Rapid and extensive transcriptional changes occur within 24 hours of infection and precede any metabolic and phenotypic changes. Within the next 48 hours, the virus activates the cells, changes their phenotypes with respect to cell size, RNA and protein content and induces metabolic pathways to cope with the increased demand for energy, supporting an efficient cell cycle entry on day three post infection. The transcriptional program that EBV initiates consists of three waves of clearly discernable clusters of cellular genes that peak on day one, two, or three and regulate RNA synthesis, metabolic pathways and cell division, respectively. Upon the onset of cell doublings on day four the cellular transcriptome appears to be completely reprogrammed to support the activated and proliferating cell, but three additional clusters of EBV regulated genes adjust the infected immune cells to fine-tune cell signaling, migration, and immune response pathways, eventually. Our study reveals that more than 98 % of the 13,000 expressed genes in B-lymphocytes are regulated upon infection demonstrating that EBV governs the entire biology of its target cell.

## Introduction

Epstein-Barr virus (EBV) was discovered in 1964 in lymphoma biopsies of patients from sub-Saharan Africa (Epstein et al., 1964; Pulvertaft, 1964). In the following years, Pope and the Henle couple independently reported that the newly termed virus induces the proliferation of immune cells from peripheral blood of any human donor (Pope, 1967; Henle, 1968). This seminal finding changed the view on the role of herpes viruses in general and provided a reliable assay for EBV’s transforming capacity mimicking its oncogenic potential. B cell transformation became the most important test to identify and investigate the contributing viral factors and to study the mechanisms leading to herpesviral latency. In the following years EBV was found to be strongly associated with other lymphomas but also gastric and nasopharyngeal carcinomas (Shannon-Lowe et al., 2017; Young et al., 2016). In vitro, EBV efficiently infects mature, resting B-lymphocytes, activates them, and induces their continuous proliferation leading to established lymphoblastoid cell lines (LCL). Many groups have contributed to the analysis of LCLs to understand the multiple viral contributions to B cell transformation and viral latency. We learnt that EBV establishes the so-called latency III program in LCLs, which includes the expression of six Epstein-Barr virus Nuclear Antigens (EBNAs) such as EBNA1, EBNA2, EBNA3A, EBNA3B, EBNA3C, and EBNA-LP, three latent membrane proteins (LMP1, LMP2A, and LMP2B), ncRNAs (EBER1, EBER2, snoRNA), and 44 miRNAs (Young and Rickinson, 2004).

Recently, the early aspects of EBV infection during the so-called pre-latent phase have gained more attention (Jochum et al., 2012b; Woellmer and Hammerschmidt, 2013), but, to our knowledge, a comprehensive analysis of the early events that lead to viral transformation of B cells is lacking. As a consequence, we do not know how the infected resting B-lymphocyte evolves to give rise to a proliferating LCL, eventually. During this initial phase of infection, the expression of certain latent genes such as EBNA2 is critical, but also lytic genes are temporarily expressed and seem to contribute to the early survival of the EBV-infected B cell promoting long-term persistence of EBV (Altmann and Hammerschmidt, 2005; Kalla and Hammerschmidt, 2012; Kalla et al., 2012). In the pre-latent phase, EBV has to overcome the initial antiviral responses of the host cell (and its organism) to be successful. Several viral processes are known that counteract these responses. The known viral processes encompass the mimicry of IL-10 by BCRF1 (Zeidler et al., 1997), repression of antigen presentation mediated by BNLF2a (Jochum et al., 2012a; Hislop et al., 2007) and certain viral miRNAs (Albanese et al., 2017) as well prevention of apoptosis by BALF1 and BHRF1 (Altmann and Hammerschmidt, 2005).

The success of infection lies in the complex strategy of EBV to reprogram the host B cell. One of the most important viral factors is EBNA2, which is supported by EBNA-LP, but several reports investigated other latent genes defining their roles immediately after infection such as the EBNA3s (Nikitin et al., 2010) or revealing the functions of the delayed expression the LMPs (Price and Luftig, 2014; Price et al., 2012; Fitzsimmons and Kelly, 2017). Initially after infection, EBNA2 plays a crucial role in cell activation and proliferation entry. EBNA2 together with EBNA-LP co-regulate the expression of important cellular genes including the proto-oncogene *MYC* (Kaiser et al., 1999) and transcription factors such as PU.1 (Zhao et al., 2011), EBF1 (Lu et al., 2016), IRF4, and CBF1 (Strobl et al., 1997; Jiang et al., 2014). These cellular genes promote B cell activation directly or activate signaling cascades, which together support cell proliferation and virus persistence during latency or prepare the latently infected cells to support the productive lytic phase including viral replication and de novo virus synthesis.

Our knowledge about EBV-mediated processes in newly infected B-lymphocytes is incomplete. Many groups have investigated the regulation and function of specific cellular and/or viral genes and processes in stable, latently infected cell lines, but the initial events that are crucial and drive the early phase of EBV infection are less known. From the perspective of the virus, EBNA-LP and EBNA2 are the two viral genes that are expressed early, but a detailed and systematic analysis of viral gene expression and their impact on host cell transcription is not available.

The aim of this study was to examine the early interactions between EBV and its cellular host. We designed time-resolved infection experiments with a focus on the B cell biology and the dynamics of cellular and viral gene regulation. Our data reveal that naïve B-lymphocytes infected with EBV undergo rapid and dramatic phenotypic changes involving cell size, content of macromolecules, metabolic and mitochondrial activities, and entry into the cell cycle. EBV imposes a strict timing of cellular genes supporting EBV’s pre-latent phase and induces the global reprogramming of the transcriptome of the quiescent, resting B-lymphocyte, which becomes an infected, activated and cycling B-blast. Our analysis suggests that the profoundest alterations at the level of the cellular transcriptome of the infected naïve B-lymphocyte occur within the first three days, whereas phenotypic and metabolic phenotypes start changing from day three onwards. Later events seem to fine-tune the biology of the host cell preparing for the ensuing phase of stable latency. The data are a rich source of cell biology covering the early molecular steps of B cell transformation driven by the tightly-controlled expression program of viral genes.

## Results

### EBV infection induces fundamental metabolic alterations in infected cells

We performed detailed time-course experiments to monitor basic metabolic parameters such as mitochondrial activity and glucose uptake in uninfected and EBV infected B cells during the first week of infection. B-lymphocytes were obtained from adenoid tissue and purified by removing cells with other identities using a negative depletion strategy (see Materials and Methods). Purified B cells were infected with the wild-type EBV strain B95-8 and a multiplicity of infection that was found optimal for infectivity and cellular survival (Pich et al., 2019; Steinbrück et al., 2015). Equivalent numbers of uninfected cells and cells infected for the indicated times (Fig. 1A,B) were stained with tetramethylrhodamine ethyl ester (TMRE), incubated with Annexin V or with 2-[N-(7-nitrobenz-2-oxa-1,3-diazol-4-yl) amino]-2-deoxy-D-glocose (2-NBDG), a glucose analogue, to analyze mitochondrial activity, early apoptosis, and glucose uptake, respectively. TMRE is a cationic red-orange dye, which accumulates in active mitochondria reflecting their membrane potential, negatively charged matrix and mitochondrial electron transport chain functions. Annexin V binds phosphatidylserine residues (PS) in the plasma membrane indicating an early phase of cell apoptosis.

**Fig. 1.**
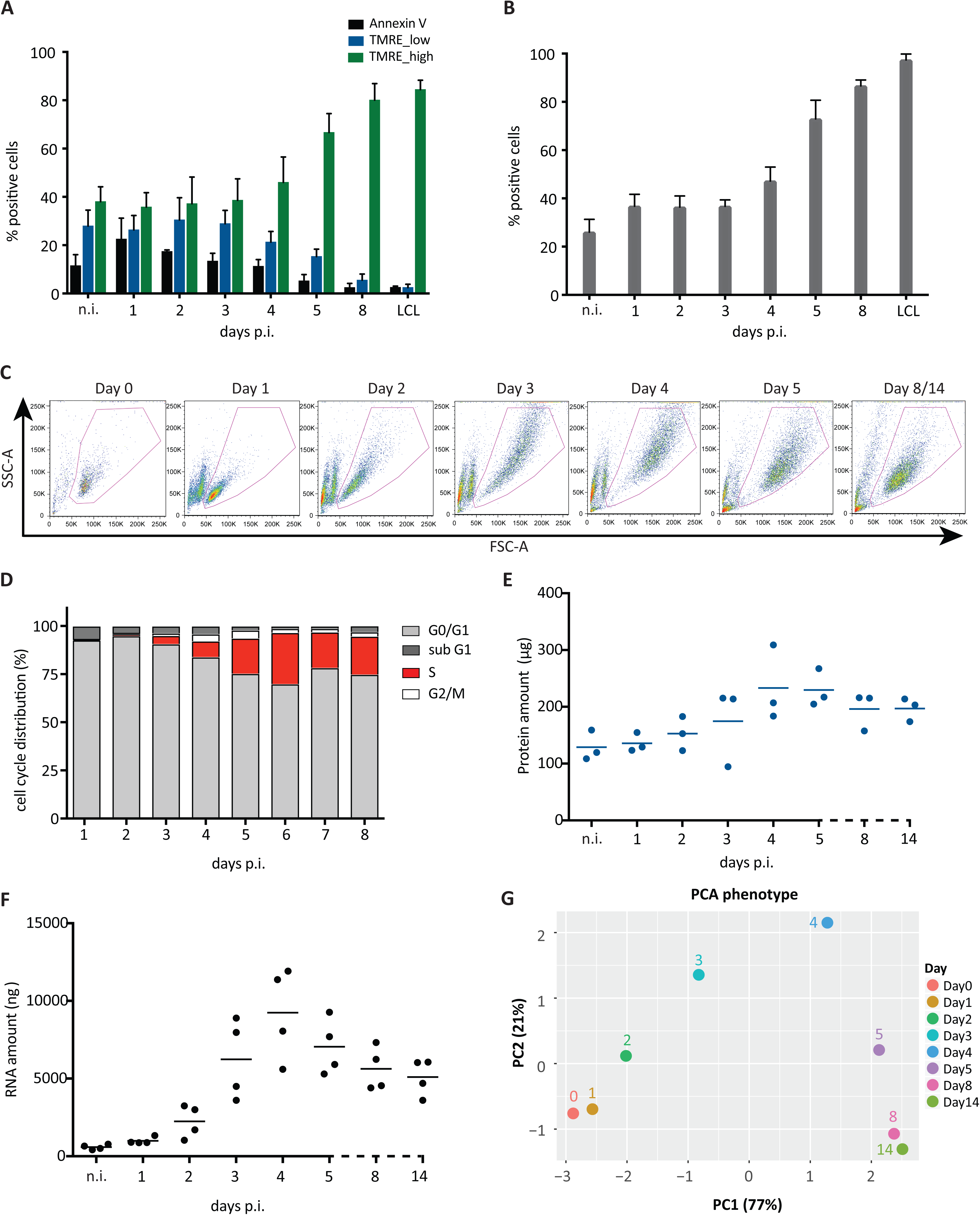
Metabolic and phenotypic parameters of B cells early after EBV infection. A. The fraction of Annexin V-positive and TMRE_low cells decreased in the course of EBV infection. The fraction of TMRE_high cells stayed constant until day three p.i., but increased starting on day four to amount to more than 80 % one week p.i.. Mean and standard deviations from four independent biological replicates are shown. B. During the first three days p.i. the fraction of cells that took up the glucose analogue (2-NBDG) was constant. Starting on day four the fraction of glucose analogue-positive cells increased up to 100 %. Bars in the graph display mean and standard deviation from four independent experiments. C. Time-resolved FACS analysis of FSC-A (x-axis) and SSC-A (y-axis) criteria of EBV-infected cells. Uninfected B cells as well as newly infected cells on day one p.i. form a homogenous population of small cells with low granularity. The cells increase in size and reach maximal size and granularity four days p.i.. Later, the cells adopt a more discrete population. On day eight and 14 post infection the cells are indistinguishable (data not shown). D. B cells after wt/EBV (2089) infection were incubated with BrdU for one hour followed by intracellular staining with a BrdU specific antibody. The cells were analyzed by FACS to determine their cell cycle distribution. Until day three the cells did not proliferate and stayed arrested in G0/G1 phase of the cell cycle. From day three p.i. the cells started to synthesize cellular DNA, as indicated by the presence of cells in the S phase. Experiments were performed using three independent replicates. E. Uninfected cells and cells infected with wt/EBV (2089) were collected daily and total protein lysates were obtained. The samples were measured using the Bradford assay. The graph shows the total amount of proteins (μg) obtained from one million cells. Measurements were performed every day after infection as indicated. The horizontal blue lines show the mean average from cells prepared from three different donors, individual samples are represented as single dots. F. Amounts of total RNA (ng) obtained from one million cells were measured using a Shimadzu MultiNA instrument. Solid black lines represent the mean average from four different donors. Each donor is represented as a single dot. G. The plot represents the principal component analysis (PCA) of the data in this figure (TMRE, 2-NBDG, cell diameter, division index, S-phase cell cycle distribution, protein content, and RNA content). The results from each experiment were normalized by the maximum value in each assay (see Methods). Next, PCA was performed and the first two components, PC1 (x-axis) and PC2 (y-axis), were plotted. The number in parentheses indicates the percentage of total variance explained by each principal component. Colored dots represent samples infected with EBV at the depicted time points.

We compared the kinetics of TMRE staining and Annexin V binding at the indicated time points prior to and after EBV infection (Fig. 1A). Two distinct TMRE-positive cell populations (TMRE_low and TMRE_high) were obvious in uninfected cells and cells infected for up to three days, which showed a similar and thus stable distribution (Supplementary Fig. S1). From day four onward, the TMRE_high population became more abundant (more than 80 % positive cells), while the TMRE_low population disappeared on day eight p.i.. The binding of Annexin V indicated a substantial fraction of apoptotic cells during the first four days of infection, which decreased starting at day five p.i. (Supplementary Fig. S1).

The significant increase of mitochondrial activity could indicate a cellular requirement for glucose, the most important source of energy. To monitor the changes in glucose uptake we analyzed uninfected and infected cells with the aid of a fluorescently labelled (FITC) glucose analogue (2-NBDG) at the indicated time points (Fig. 1B). Cells were transferred to low glucose medium and incubated with 100 μM 2-NBDG for one hour under optimal culture conditions. The accumulation of the 2-NBDG fluorescent signal in the cells was measured by FACS over time. Similar to the TMRE results, the uptake of glucose did not change in infected cells compared with uninfected B cells up to three days p.i. with levels of about 40 % 2-NBDG positive cells (Fig. 1B; Supplementary Fig. S2). Four to five days p.i. the uptake of the glucose analogue increased substantially and almost all cells were found to become 2-NBDG positive, eventually.

Based on these results we conclude that mitochondrial activity and glucose uptake increase in the time course of infection. Moreover, within the first three days p.i. the metabolism of the infected B cells is comparable with uninfected, resting B-lymphocytes, but starting from day four onward, the infected cells increase their metabolic rate which is maintained in established, stably proliferating LCLs.

### Phenotypic changes in the host B cells during early time points after virus infection

Next to the basic metabolic parameters we studied additional cellular phenotypes of the EBV infected cells. Quiescent naïve B-lymphocytes (IgD^+^, CD38^-^) were purified from adenoid tissue by physical sorting. Visual, daily inspection of the infected cells (Fig. 1C) demonstrated that they grew massively in size and gained granularity according to forward (FSC-A) and sideward scatter (SSC-A) FACS criteria, respectively. The population of cells started to become more homogenous on day five p.i. to reach a state that was maintained in established lymphoblastoid cell lines (Fig. 1C). We determined the cells’ diameter during EBV infection (Supplementary Fig. S3A). Initially, the diameter of the purified naïve B cells was found to be 5.5 μm. The cells grew constantly up to 9 μm on average within the first four days of infection, but their diameter shrank to 8 μm later and did not change further when a stable population of lymphoblastoid cells emerged, eventually (Supplementary Fig. S3A).

EBV infection of the resting naïve B lymphocytes induced their delayed entry into the cell cycle. Naïve B cells were found arrested in G0/G1 for two days post infection according to their DNA content and metabolic incorporation of BrdU (Fig. 1D). The first cells entered S phase on day three p.i. and the fraction of S phase cells increased on the following days. The B cells underwent substantial DNA synthesis as indicated by the fraction of cells in S-phase on day five and six p.i. exceeding this number in established latently infected lymphoblast populations. In parallel to the analysis of the cell cycle we monitored the dynamics of cell divisions by loading the uninfected B-lymphocytes with CellTrace Violet and by following its distribution by FACS to calculate the division index (i.e. the average number of divisions for all cells, Roederer, 2011) in the course of EBV infection (Supplementary Fig. S3B). Within the first three days the infected B cells did not proliferate. The first cell division was observed on day four p.i., immediately followed by a two-day long phase with a steep increase of the division index indicating a generation time of only 12 hours. From day seven p.i. onward, the cells decelerated and divided once within a period of 24-36 hours.

The EBV-induced B cell growth and cellular proliferation resulted in global alterations of macromolecules in the infected cells. The contents of total cellular RNA and protein in 1×10^6^ intact cells that had been physically sorted according to their forward and sideward scatter criteria were measured daily and compared with the lymphoblast cell population at two weeks p.i. (Fig. 1E,F). In cells infected with EBV for four days the total protein content was about 2.5-fold higher compared with uninfected cells (Fig. 1E). At later time points the protein content decreased slightly and became stable at a higher level than in resting B-lymphocytes after about one week p.i.. Similarly, the total cellular RNA content increased substantially by a factor of fifteen when it peaked at day four p.i. (Fig. 1F). A single non-infected B-lymphocyte contained about 0.59 pg RNA on average, whereas an EBV-infected activated B cell contained about 9.2 pg on day four p.i.. Starting from day five, the amount of single cell RNA decreased constantly and stabilized at about 5.1 pg a fortnight p.i. (Fig. 1F).

The results from the metabolic and phenotypic assays were normalized (see Materials and Methods) and combined via Principal Component Analysis (PCA) to visualize the global dynamics of the metabolic and phenotypic changes during EBV infection. The PCA shows that uninfected and infected cells were similar in terms of their metabolic functions and phenotypes during the first two days p.i. (Fig. 1; Supplementary Fig. S3) since they cluster close to each other (Fig. 1G). At day three and four p.i. the cells differed substantially from the initial cluster (Fig. 1G) due to their increased energy consumption, entry into the cell cycle, and first cell divisions (Fig. 1A,B, and D; Supplementary Fig. S3). Infected cells on day 5 p.i. differed even more from cells during the previous days of infection, but they were closer to cells infected for eight and 14 days demonstrating their similar metabolic processes and phenotypes (Fig. 1G). Overall, this analysis revealed that the changes in metabolic and phenotypic features are very limited within the first two days of infections, whereas they massively change between day 2 and day 5 p.i. Already on day 8 p.i. the cells reached a stable, latent cell phenotype, which reflects their established and final cellular fate.

### EBV infection causes global alterations in the B cell transcriptome

The prominent phenotypic changes of naïve B cells early after EBV infection indicated comprehensive alterations in the entire B cell biology. To analyze the detailed dynamics of gene expression in the course of infection we performed sequential, time-resolved RNA-seq experiments with non-infected primary B-lymphocytes and with cells infected with EBV from day one to day five, daily, and on day eight and on day 14 p.i.. Sorted, quiescent naïve B cells (IgD^+^/CD38^-^) from three different donors were infected with the wt/B95.8 (2089) EBV strain (Delecluse et al., 1998) with an optimal multiplicity of infection (Steinbrück et al., 2015; Pich et al., 2019). Each day post infection 10^6^ living, physically intact cells were collected by FACS sorting according to their forward and sideward scatter criteria. We extracted their RNA and established polyA enriched libraries to allow subsequent whole transcriptome sequencing on an Illumina HiSeq 4000 instrument with a high coverage. After gene counting but prior to any downstream analyses, we performed quality controls to ensure that all libraries were of acceptable quality. All but one sample from day 8 p.i. passed the quality controls (Supplementary Fig. S4). Supplementary Figure S5 provides the chart of the entire bioinformatic workflow and detailed information can be found in Materials and Methods.

Initially, we focused on the global transcriptional changes during infection (Fig. 2). A PCA was performed on the log-transformed gene expression matrix. The first two principal components, representing the two directions of largest variance (PC1 and PC2) are shown in Figure 2A and the expression levels of the top 100 genes that contributed the most to PC1 and PC2 are displayed in heat maps in Supplementary Figure S6.

**Fig. 2.**
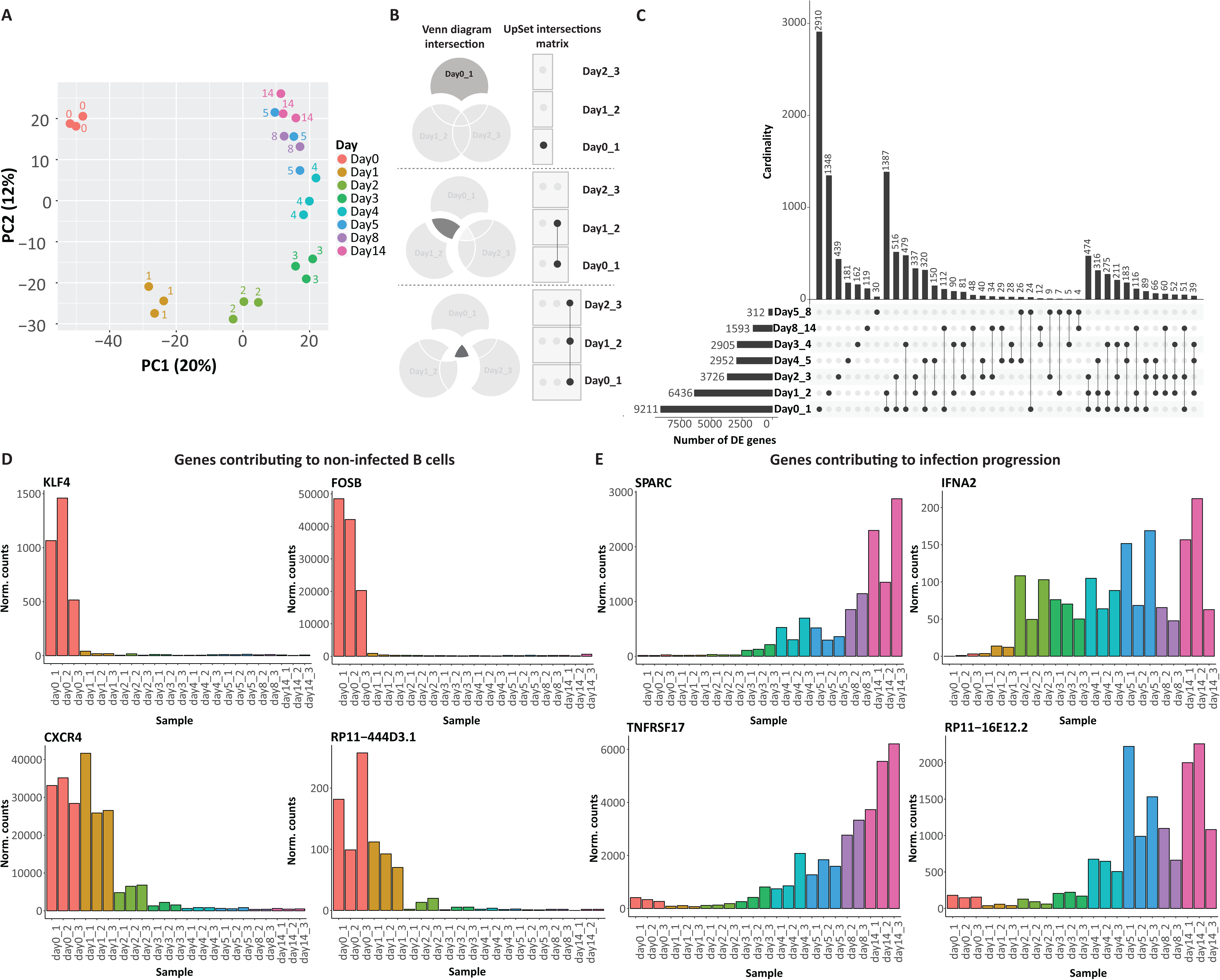
Analysis of the time-resolved RNA-seq data. A. Principal Component Analysis (PCA) of the samples from sequencing libraries at the given time points post infection (p.i.). The daily samples are represented by three biological replicates with the exception of day 8 (n=2). The samples are shown as a function of the first (PC1) and the second (PC2) principal components. The x and y-axes show the percentages of variance explained by PC1 and PC2. B. The three Venn diagrams serve as examples to illustrate the syntax of the UpSet diagram shown in panel C. The single dot and dots with interconnecting vertical lines form the intersection matrix. Paired intersections are depicted as black dots, grey dots indicate the sets that are not part of the intersection. Black lines connecting two or more black dots indicate which sets form the intersections. C. The UpSet plot visualizes intersecting sets of differentially expressed (DE) genes at different time points after EBV infection. Groups of DE genes, obtained from the pairwise comparison (performed by DESeq2; see Methods) as indicated (Day0_1 = day 0 versus day 1, etc.) form seven sets in the UpSet matrix, which are depicted as black horizontal bars with “Number of DE genes” to indicate the set sizes. Each row represents a matrix set. The columns show the intersections depicted as areas in the schematic Venn diagram as explained in panel B. The heights of the black columns indicate the cardinality (number of elements) in the different intersections. D. The bar plots show the time-resolved expression of four selected genes, which displayed high expression in non-infected cells and/or only in cells very early after EBV infection. The four genes are among those with the highest loadings on PC2. The x-axis displays samples from different days and three different donors, whereas y-axis depicts normalized read counts. E. Examples of four genes that contributed most to PC1. The indicated genes were barely expressed in uninfected cells but became activated at different time points post EBV infection.

First, this analysis showed that the three biological replicates of each time point had a very low variability, since they were very close to each other in the PCA plot. Further, the PCA clearly separated the samples obtained from uninfected cells and infected cells of the first three days, which formed four separate groups (Fig. 2A). In contrast, samples from day four onward were close together. This observation suggests that the profoundest variations in gene expression occurred already during the first three days of infection in contrast to the phenotypic and metabolic changes that primarily occurred on day three and later (see Fig. 1). After day 3 the transcriptional changes were more limited and probably restricted to specific cellular processes.

This observation was confirmed when looking at the DE genes between consecutive days after EBV infection, which we visualized using the Upset package (Lex et al., 2014) (Fig. 2B and C). The most abundant group of DE genes (n=9,211) was identified comparing uninfected B cells and cells infected for 24 hours, and the second most abundant group of DE genes (n=6436) was found when comparing Day1 with Day2. The pairwise comparison between the following days demonstrated a constant decrease in the numbers of DE genes over time (Fig. 2C; Supplementary Fig. S7).

Similarly, the set analysis also uncovered a gradual time-dependent reduction in the sizes of the sets (i.e. the cardinality or set size describing the number of genes per set) that belonged to unique subsets or set interactions (Fig. 2C). This finding indicated that the number of DE genes decreased substantially at later time points (e.g. Day4_5, Day5_8, and Day8_14), when latency was established. For instance, 2,910 genes changed specifically between day 0 and day 1, while only 119 genes were differentially expressed comparing the last two time points (day 8 and day 14).

Having identified sets, subsets and their multiple intersections in a time-resolved approach we analyzed the expression kinetics of potentially interesting genes selected among those that explain most of the variation along the first two principal components (Supplementary Fig. S6) and might be functionally relevant in the course of viral infection (Fig. 2D and E).

Four genes, *CXCR4,* a chemokine receptor, two transcription factors *(KLF4* and *FOSB),* and ncRNA *(RP11-444D3.1)* were highly expressed in uninfected cells but their expression was lost within 24 *(KLF4* and *FOSB)* or 48 hours p.i.. Interestingly, *CXCR4,* which is involved in hematopoiesis and migration of resting leukocytes (Sugiyama et al., 2006), was downregulated after B cell activation mediated by CD63 and IL-21 (Yoshida et al., 2011). *FOSB* is a proto-oncogene and G0/G1 switch protein, which is induced upon B-cell receptor induction (Yin et al., 2008). The transcription factor KLF4 is involved in a G0/G1 cell cycle arrest, reduces B cell proliferation and was also identified to be downregulated after B cell activation by CD40 (Yusuf et al., 2008; Klaewsongkram et al., 2007). The functional significance of these four cellular genes (Fig. 2D), which exemplify a large gene set is speculative, but they might contribute to the quiescent state of naïve B-lymphocytes, which EBV overcomes.

In contrast, the expression of a different group of genes gradually increased during the progression of EBV infection. This is exemplified by *SPARC,* Interferon alpha 2 (*IFNA2*), TNF Receptor Superfamily 17 (*TNFRSF17*), and the ncRNA (*RP11-16E12.2)* (Fig. 2E). SPARC encodes a glycoprotein, which has emerged to correlate with progression of many cancers, is involved in extracellular matrix synthesis and promotes changes of the cell shape (Sangaletti et al., 2014; Chen et al., 2012). *IFNA2* is a cytokine and interferon secreted by cells in response to viral infection (Fensterl et al., 2015; Gresser, 2015). *TNFRSF17,* known also as B-cell maturation antigen (*BCMA)* was found to be induced in Burkitt lymphoma cells after *RUNX3* upregulation, which plays an important role in the proliferation of EBV-infected human B cells (Brady et al., 2009). No function has been ascribed to the ncRNA *RP11-16E12.2.* The expression of these four genes that exemplify a large gene set was induced about three to four days post infection and increased further with time.

In summary, this analysis revealed global changes in transcription that EBV induces in naïve primary B-lymphocytes. The most dramatic transcriptional changes occurred during the first three days of infection affecting the entire biology of the host cell and paving the way to the profound phenotypic and metabolic changes we observed starting from day 3 onwards (Fig. 1).

### Gene expression cluster analysis identifies multiple and specific biological processes regulated by EBV infection

Next, we wanted to exploit our time-resolved dataset to classify genes according to their expression dynamics, aiming to pinpoint specific biological processes that EBV drives at the different time points post infection. To this aim, we followed the computational strategy illustrated in Figure 3. First, we selected all genes with a dynamic behavior, defined as the union of all genes significantly differentially expressed (DE) (FDR<0.1) with a fold-change greater than 2 between at least two time points (see Methods). We identified 11,178 genes that satisfy these criteria, representing 86% of all genes tested (n=13,007). On the other hand, only 241 genes (1.85% of all tested genes) did not show any significant change in their expression levels during the course of EBV infection. This finding suggests that EBV globally affects the entire transcriptome in the pre-latent phase of infection.

**Fig. 3.**
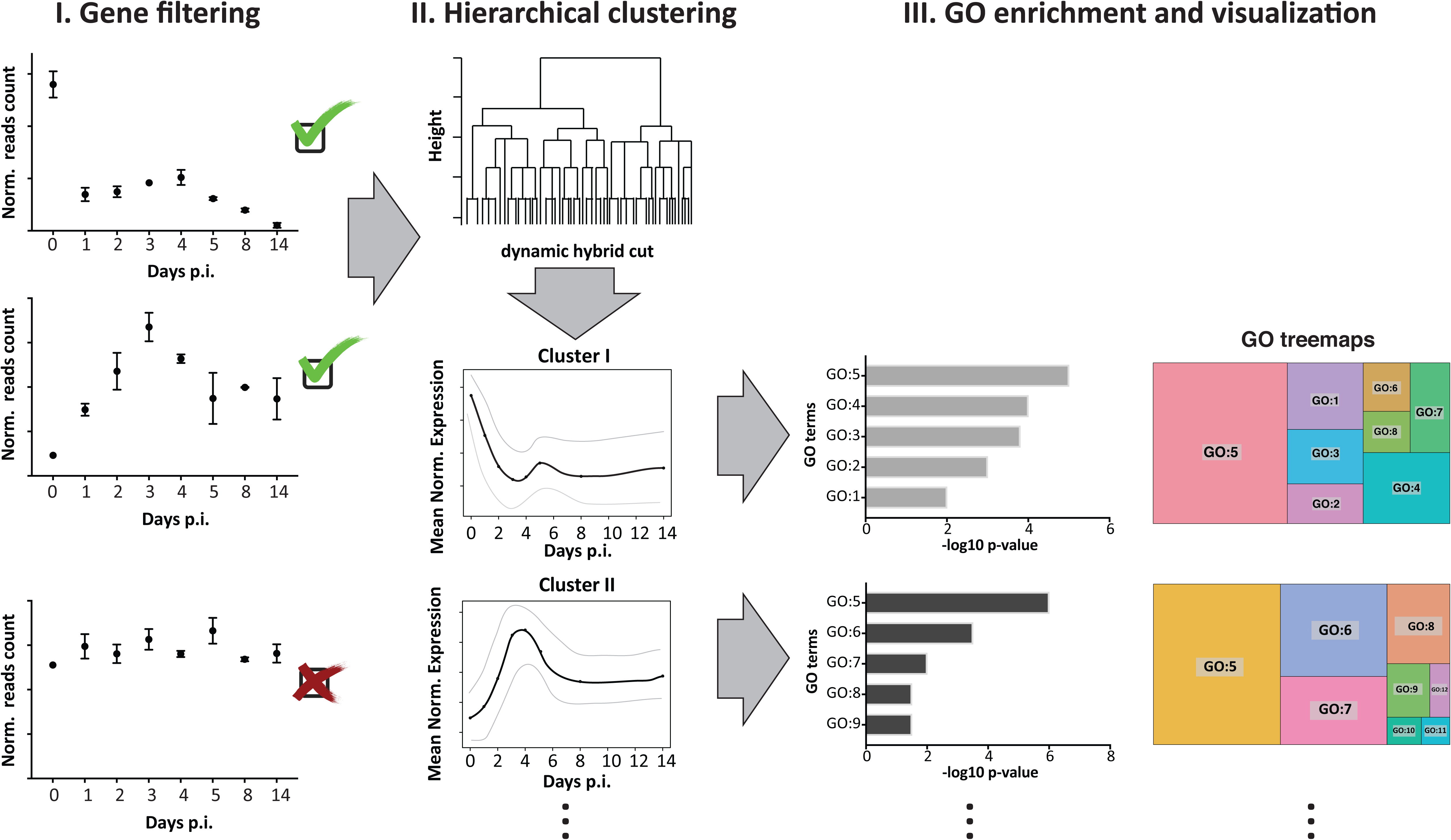
Workflow of the bioinformatic analysis and gene clustering. The schematic overview represents the three main steps of the bioinformatic analysis: gene filtering, hierarchical clustering, and GO enrichment and visualization performed to identify specific clusters of differentially expressed genes and related biological processes.

The significant DE genes were classified into six robust clusters (Supplementary Fig. S8), each characterized by different expression dynamics (Fig. 4; see Methods for details on gene filtering and clustering). Finally, a gene ontology (GO) enrichment analysis identified biological processes within each cluster of genes (Fig. 4; Supplementary Fig. S9).

**Fig. 4.**
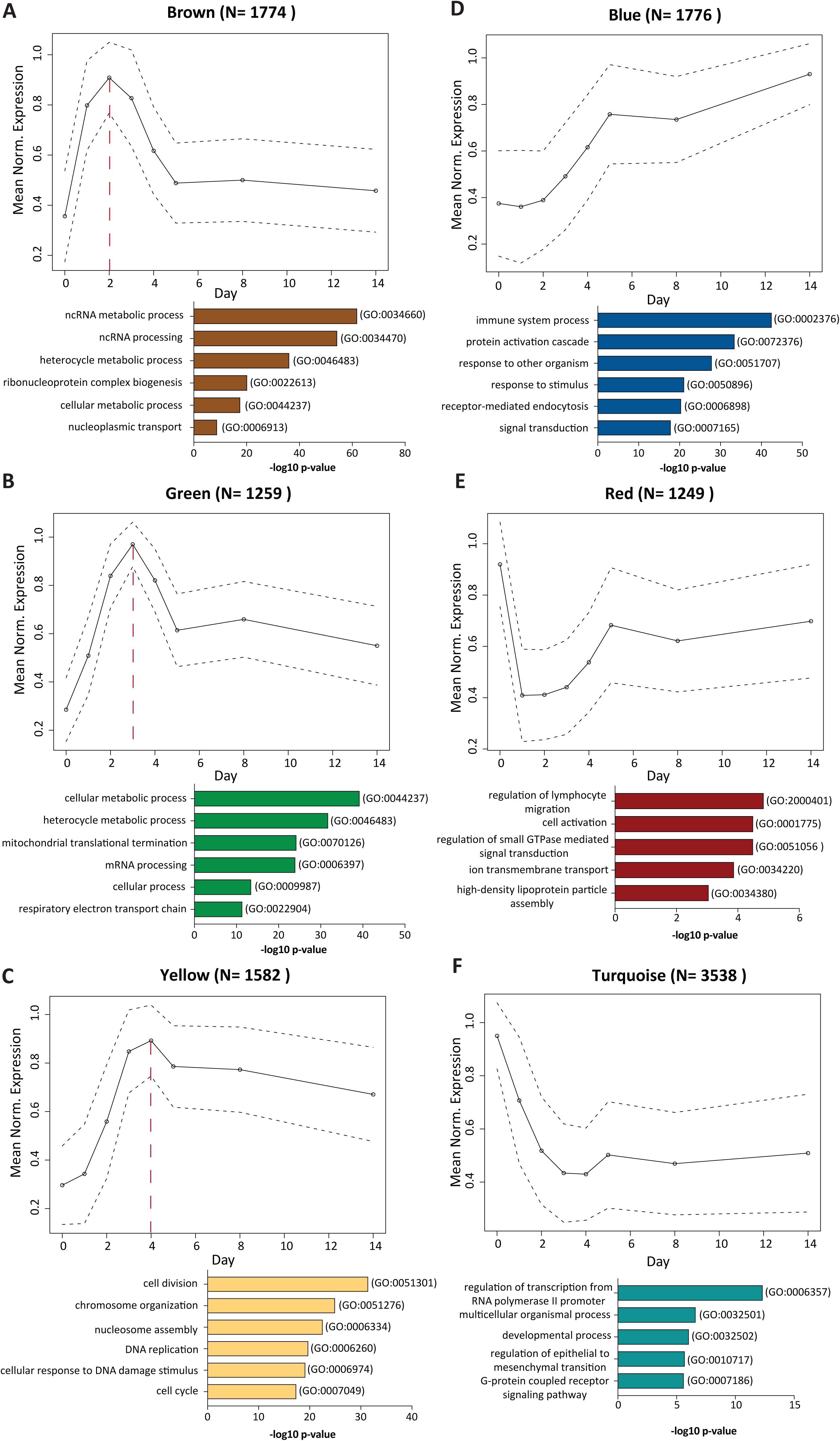
Cluster analyses of differentially expressed (DE) genes in the course of EBV infection define six clusters of genes with specific expression dynamics and the corresponding biological processes. Six clusters of genes are shown with their arbitrarily assigned color name codes and mean expression levels. The y-axis represents Mean Normalized Expression as a function of time (days p.i.) on the x-axis. The numbers on the top of the plots (N) indicate the numbers of the genes that form the cluster. The circles indicate average expression levels, while the dashed lines delineate the standard deviations. The red, dashed vertical lines mark the expression peak of three clusters. All genes within a given specific cluster were used to perform a gene ontology (GO) analysis that is summarized in a bar plot below the depicted graphs. The plots contain the top 5 or 6 enriched GO terms with their GO specific IDs (on the right) according to their –log10 p-value (x-axis). The color of the bars relates to the color name of the clusters. A. The graph depicts the expression pathway of the brown cluster. This group contains genes that peaked on day two p.i., decreased until day five and showed stable levels thereafter. The GO analysis identified the most significant biological processes involved in ncRNAs metabolism and processing, heterocycle metabolism, and ribonucleoprotein complex biogenesis. B. The green cluster peaked on day three p.i.. Within this cluster, the most frequently found GO terms were cellular metabolic process and mitochondrial biogenesis such as mitochondrial translational termination or mitochondrial processes linked to energy production as the respiratory electron transport chain. C. Genes in the yellow cluster peaked on day four after EBV infection and decreased slightly thereafter. The analysis pointed to the enrichment of biological processes categorized into cell division, chromosome organization, DNA replication, and cell cycle processes. D. The blue cluster encompasses genes that constantly but slowly increased during infection. The majority of these identified genes belong to the immune system process, protein activation cascade or are genes that are involved in response to external stimulus. E. The group of genes in the red cluster is characterized by an initial loss of expression on the first day after infection and a rapid recovery until day five. The GO analysis of this cluster highlighted biological processes related to lymphocyte migration, cell activation, or regulation of small GTPase mediated signal transduction. F. The graph displays pathways present in the turquoise cluster. It encompasses genes that are highly expressed in uninfected cells but downregulated in infected cells until day four p.i. when the genes adapted a reduced but stable expression level. GO terms enriched in this group of genes indicated processes of cell development and multicellular organismal process. In this group of genes, very broad GO terms were found.

In particular, we distinguished two groups of clusters: three clusters had well-defined individual peaks on day 2, 3, or 4 p.i. (Fig. 4A,B, and C). In contrast, three clusters had peaks either in non-infected cells or two weeks after infection (Fig. 4D,E, and F).

The genes whose expression levels peaked on day 2, 3, or 4 were strongly enriched for very specific GO terms (Fig. 4A,B,C; Supplementary Fig. S9). In particular, the brown cluster (Fig. 4A) displayed the peak of expression on day two p.i. and is enriched with genes involved in mostly ncRNA metabolism and processing, ribonucleoprotein complex biogenesis, or nucleoplasmic transport. The green cluster with its peak of gene expression on day three p.i. (Fig. 4B), included genes that are involved in mitochondrial translation termination, the respiratory electron transport chain, and cellular metabolism, in agreement with our metabolic data in Figure 1A and B. Third in the group, the yellow cluster (Fig. 4C) had its peak of expression on day four p.i., when the infected cells start to proliferate and undergo the first cell divisions (Fig. 1D; Supplementary Fig. S3B). This cluster with GO terms related to chromosome organization, cell cycle process, cell division, and DNA replication faithfully reflects the phenotypic changes in cell biology we recorded in our initial set of experiments (Fig. 1).

The second group of clusters (Fig. 4D,E,F) included the blue cluster (Fig. 4D), which showed a constant increase in gene expression during the course of EBV infection. This cluster is enriched for discrete biological GO processes including the immune system process, the protein activation cascade, which consists of a sequential series of enzymatic modifications (proteolysis, covalent modifications, binding events, etc.), and defense responses to other organisms including viruses. On the contrary, the red cluster (Fig. 4E) was characterized by an immediate decrease of cellular gene expression followed by a rapid increase in expression levels and included genes mostly involved in lymphocyte migration, cellular activation, and regulation of GTPase activity. Additionally, the turquoise cluster (Fig. 4F) displayed an immediate loss early after infection and a significant stable reduction of gene expression throughout the observation period. This group of genes was enriched in GO categories including transcription from RNA polymerase II promoters as well as developmental processes, which might reflect the final status of lymphoblast cells that do not further differentiate *in vitro.*

The cluster analysis defined very specific dynamic patterns of a plethora of biological processes that are regulated during the progression of EBV infection. Some of these processes are related to the metabolic and phenotypic changes that we observed, confirming the results from experiments shown in Figure 1 and providing a list of specific genes associated to those changes (green and yellow cluster). Moreover, additional processes not previously linked to EBV infection (e. g. ncRNA processing, mitochondrial activity, or a group of various genes that decrease their expression immediately after infection) were identified and their dynamics was characterized. Within a very strict and well-defined time window, the virus completely changes the transcriptome of resting, naïve B cells, which is followed by specific metabolic and phenotypic changes (Fig. 1). The most prominent transcriptional alterations take place very early during the first three days of infection, when many cellular processes are induced or downregulated. Later during infection cellular genes undergo relatively minor changes to adopt the stable gene expression profile of the activated and EBV transformed, immortalized B cells.

### Regulation of viral genes expression in the early phase of EBV infection

In parallel with cellular genes that are involved in cellular activation and transformation, viral genes are expected to show a very dynamic but regulated expression during the pre-latent early phase of EBV infection. We knew from published work that initially after infection EBV latent genes are induced (Jochum et al., 2012b; Price and Luftig, 2014; Kalla and Hammerschmidt, 2012) but also certain lytic genes are expressed albeit only temporarily. Some of the latter have essential or auxiliary roles supporting the early survival of the infected B cells (Altmann and Hammerschmidt, 2005; Kalla et al., 2010; Wen et al., 2007).

Similar to the study of cellular genes, our transcriptomic data also allowed a very detailed analysis of viral genes and their regulation. We found that the relative abundance of viral genes with respect to cellular genes peaked immediately after infection but started to decline within 48 hours (Fig. 5A). Viral gene expression at the early time points after infection also implicated differences defined with PCA of viral genes (Supplementary Fig. S10A). The samples from day one and two differed the most, and starting from day three the samples formed a more homogenous group indicating their similarities in terms of viral gene expression. However, different viral genes showed different dynamics. In our RNA-seq experiments followed by hierarchical clustering (Fig. 5B; Supplementary Fig. S10B), we identified three clusters with viral genes that showed specific gene expression dynamics and kinetics during the pre-latent phase of infection (Fig. 5C,D,E; Supplementary Fig. S10B). Certain viral genes in cluster I, including EBNA2/BYRF1, EBNA-LP, and BHRF1 (Fig. 5C), are characterized by an extraordinarily high expression, especially during the first two days p.i.. These genes reached their highest expression levels on day one, but they declined substantially by more than a factor of three within the next 48 hours (Fig. 5C). Other genes in cluster I showed a very similar expression dynamics, but at significantly lower levels (Supplementary Fig. S11). The expression levels of viral genes in cluster II peak around day 2 and 3, including genes such as EBNA3C or BKRF1/EBNA1 (Fig. 5D), which are known to follow the initial EBNA2 expression. In contrast to the majority of DE viral genes, the cluster III contained viral latent genes such as BNLF1/LMP1 and LMP2A (Fig. 5E), which accelerated their expression starting from day two after EBV infection reaching modest, often variable levels of expression two weeks post infection (Supplementary Fig. S11).

**Fig. 5.**
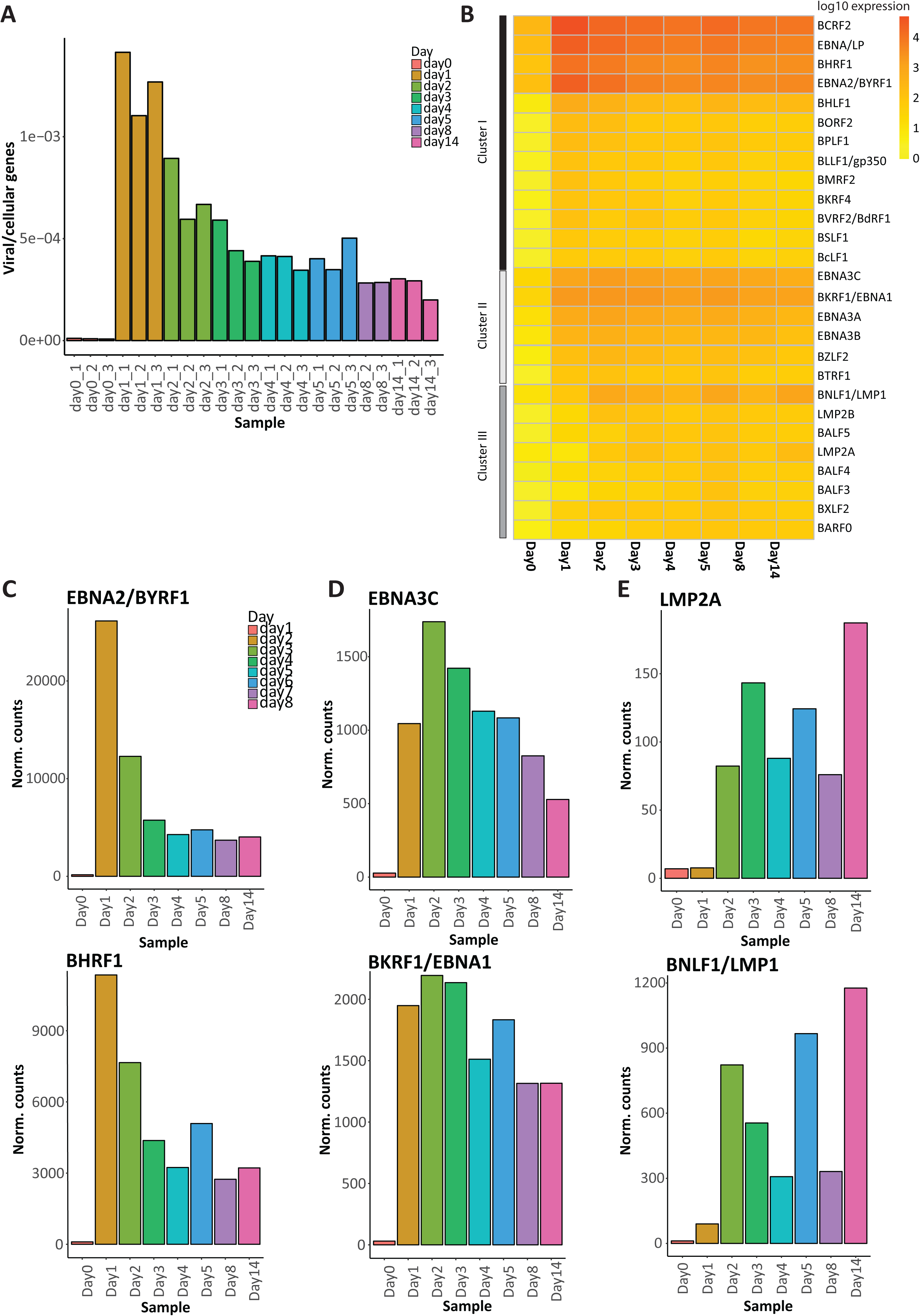
Viral gene expression during the course of early EBV infection and activation of naYve B cells. A. The ratios between counts allocated to viral genes versus counts allocated to cellular genes were calculated at different time points p.i.. As expected, no viral genes were present in the uninfected, naïve B cells (Day 0). On day one, the relative abundance of viral genes peaked but decreased subsequently during the process of EBV infection and establishment of latency. B. Heat map of normalized log10-mean expression values of significantly DE viral genes. Colors from yellow to red indicate the range of calculated values from low expression level to high expression level, respectively. Three clusters I, II, and III are indicated on the left and refer to panels C, D, or E, respectively. C. Two examples of viral genes of cluster I, EBNA2/BYRF1 and BHRF1, during the course of EBV infection. The x-axis depicts different days p.i. and the y-axis indicates the mean of normalized counts from three different biological replicates. This group of viral genes has its peak expression on the very first day p.i. and declined constantly thereafter. D. Expression plots of two examples of viral genes found in the cluster II, EBNA3C and BKRF1/EBNA1, at defined time points after EBV infection. The genes’ peak of expression is on day two or three p.i. followed by a continuous decrease of expression. E. Expression plot of two latent viral genes, LMP2A and BNLF1/LMP1 included in cluster III. The expression of these genes increases over time post EBV infection.

The most dynamic regulation of EBV gene expression took place during the first two days after infection (Supplementary Fig. S10A) pointing to a very stringent expression program in the pre-latent phase. The specific expression patterns of viral genes strongly correlated with the most prominent changes in the host B cells (Fig. 1 and Fig. 2A), consistent with viral genes being the main trigger of B cell activation and transformation in vitro.

To summarize, we provided a detailed, time resolved analysis of the transcriptome of B cells during the EBV infection, looking at both cellular and viral genes. All our processed data, including the lists of genes in each cluster, are available via a user-friendly R Shiny app (https://scialdonelab.shinyapps.io/EBVB/) that provides an easy interface to explore cellular and viral genes and their regulation in the pre-latent phase of EBV infection.

## Discussion

### A detailed, time resolved characterization of the transcriptome of infected B cells

The dramatic metabolic and phenotypical changes we saw in our experiments (Fig. 1; Supplementary Figs. 1-3) raised questions about the transcriptional events driving them. To address these questions, we performed a time-resolved RNA-seq experiment at an unprecedented resolution: we collected and processed samples daily during the first five days of infection, and then at day 8 and day 14 post infection. After removing technical (e.g. sequencing depth) as well as biological (e.g. RNA content) confounding effects through data normalization, we devised and applied a computational strategy consisting of several steps (see Fig. 3 and Supplementary Fig. S5) that allowed us to: (i) detect genes that change their expression levels during EBV infection; (ii) define clusters of genes sharing similar expression patterns; (iii) characterize the different steps of the B cell reprogramming by identifying the biological processes involved and their dynamics.

We observed rapid and extensive transcriptional changes that occurred already within 24 hours of infection and preceded any metabolic and phenotypical changes. Moreover, we found new processes (e.g. ncRNA processing) that have not been previously linked to EBV infection. We also analyzed the dynamics of viral genes, that are dominated by the immediate maximal expression of three viral genes (EBNA2, EBNA-LP, and BHRF1) already on day 1 post infection

Our time-resolved transcriptional dataset represents an important resource for the entire community. Hence, to facilitate access to it, we created a website (https://scialdonelab.shinyapps.io/EBVB/) where the processed data can be downloaded and explored: via an easy web-based interface the dynamics of all cellular and viral genes can be tracked starting from uninfected B cells, through the pre-latent phase of EBV infection, until the establishment of LCLs two weeks post infection.

Overall, our data indicate that EBV triggers a plethora of substantial changes in the biology of the entire host cell. In a well-defined and short interval of time, the virus drastically overturns the transcriptional program of resting and quiescent B cells and reprograms a multitude of different biological processes and pathways.

### B and T cells differ in their activation kinetics

Alterations in the metabolism of immune cells, collectively termed immunometabolism (Murray et al., 2015) trigger important changes in lymphocytes including their activation, proliferation, and eventual differentiation into different subtypes of multipotent precursor, effector or memory cells. After antigen encounter naïve and quiescent T cells undergo activation and start cell divisions within 24 h, which is in stark contrast to B-lymphocytes infected with EBV as demonstrated in this paper. We also compared our infection model with a physiological stimulus, CD40 ligand and IL-4, that together activate and induce long-term cell divisions of primary B-lymphocytes (Banchereau et al., 1991; Wiesner et al., 2008). We found that this stimulation recapitulates all aspects of EBV infection (Pich et al., 2019) suggesting that EBV does not inaugurate a special program of cellular activation and cell cycle entry but rather usurps physiological circuits. The differences between the two classes of B and T lymphocytes might stem from the fact that T cells are programmed to react immediately upon antigen encounter, become activated within minutes and start dividing every four to six hours (van Stipdonk et al., 2003). To support this acute process, the lymphocytes have to adapt their metabolism in a very short time to fulfill the needs of increased energy and biomolecules in different microenvironments and different parts of the entire organism. These dynamic changes in basic bioenergetic processes enable the cells to modulate their catalytic and anabolic reactions in a very rapid and effective manner (Slack et al., 2015).

Upon activation, the naïve T cells rely mostly on OXPHOS production of energy to realize the basic need of energy to promote cell survival. The activated T cells are characterized by an elevated glycolysis and glutaminolysis, which provide the cells not only with ATP but also with the most important precursors for newly made biomolecules such as nucleic acids, amino acids, lipids etc. (Wang and Green, 2012; Buck et al., 2015; Almeida et al., 2016). Glucose as a primary fuel has an important role during T cell activation as well as subsequent differentiation since blocking of glucose catabolism inhibits T effector cell function *in vitro* and *in vivo.* Activated CD8^+^ T cells shift from fatty acid oxidation to aerobic glycolysis and glutamine oxidation to support their differentiation into cytotoxic T cells and generation of memory CD8+ T cells (Cao et al., 2014).

Similarly, upon CD40/IL-4 stimulation B cells synthesize and accumulate macromolecules and gain in volume. B-cell receptor (BCR) crosslinking as well as IL-4 and TLR-mediated activation increase the uptake of glucose and aerobic glycolysis concomitant with the upregulation of glycolytic enzymes (Doughty et al., 2006; Dufort et al., 2007). Infection of primary B cells with EBV leads to cell activation, which mimics the physiological signaling such as BCR signaling pathway mediated by LMP2A (Mancao and Hammerschmidt, 2007) or CD40 stimulation supported by LMP1 (Kieser and Sterz, 2015). Our results confirm that EBV infection also causes substantial changes in cell growth (Fig. 2A and B), cell cycle (Fig. 2C and D), and cellular content of macromolecules such as protein and RNA (Fig. 2E and F). Activating adjustments in the cellular phenotype increase the demand for energy and biomolecules to support cellular functions and cell divisions. The uptake of glucose increased after a three day long latency period and started on day four p.i. (Fig. 1B) indicating the upregulation of glycolysis only prior to the highly proliferative stage of the infected B cells on day five and six p.i.. Thus, the increase of glucose uptake and the activation of immunometabolic functions are late events in B cells induced by EBV infection, which might not differ from physiological activating signals such as antigen encounter of these immune cells in vivo.

In addition to glucose metabolism, mitochondria are key actors in modulating bioenergetic (catabolic and anabolic) processes in the cell by providing ATP but also ROS and precursors for the synthesis of other molecules. T cell activation supports mitochondrial activation and biogenesis (Ron-Harel et al., 2015, 2016). The generation of mitochondrial ROS activates the important nuclear factor of activated T cells (NFAT) and promotes the release of cytokines (Sena et al., 2013; Weinberg et al., 2015). In our infection experiment, mitochondrial activity increased starting from day four p.i. (Fig. 1A), when the cells enter the cell cycle, replicate DNA and start dividing (Fig. 2C and D).

T cell activation induces many different signaling pathways that control diverse metabolism-associated factors such as Myc, HIF-1α, mTOR, or Gsk3. The proto-oncogene Myc is known to upregulate glucose, glutamine, and polyamine metabolism during T cell activation, similar to the upregulated metabolic pathways in cancer cells (Rathmell, 2011; Wang et al., 2011; Stine et al., 2015). Antigen-specific immune responses by B cells also affect their metabolic program and activate HIF-1α and Myc. Especially Myc is crucial for the induction of the glucose transporter (GLUT1) during B cell activation (Cao et al., 2014) and enhances the expression of hexokinase 2 (HK2) and pyruvate dehydrogenase kinase-1 in Burkitt lymphoma cells (Kim et al., 2008). Oncogenic Myc levels increase the mitochondrial mass and oxygen consumption since several Myc targets are involved in mitochondrial biogenesis and functions (Kim et al., 2008; Scarpulla, 2008). The main driver of B cell transformation during EBV infection is EBNA2, which directly targets *MYC* and promote its expression (Kaiser et al., 1999; Liang et al., 2016). In fact, transcript levels of EBNA2 reach maximal levels on day 1 after EBV infection followed by extreme levels of Myc on the next day (this study and Pich et al., 2019). Thus, very early and EBNA2 induced high levels of Myc seem to be the main trigger for B cell activation and reprogramming in the pre-latent phase of EBV infected cells.

### Dynamic and comprehensive adjustments in the host cell transcriptome early after infection

To our knowledge, only few publications describe global cellular changes after viral infection focusing on the expression and regulation of protein-coding as well as ncRNA genes. A recent report studied the host transcriptome and proteome during KSHV infection and identified global changes in the cellular networks that are essential for virus latency (Sychev et al., 2017). The authors identified alterations in approximately 17 % of the host transcriptome and 13 % of the cellular proteome that includes also dramatic changes in the phosphoproteome 48 hours after KSHV infection. EBV is infecting a cell not yet in the proliferation cycle whereas Sychev et al. in their KSHV infection model used Tert-immortalized, proliferating microvascular endothelial cells. Similar to KSHV, EBV establishes a latent infection, but the impact on the host cells is much more dramatic. EBV reprograms the B cells and we found statistically significant changes for all but 241 cellular genes corresponding to 1.85 % of all 13,007 tested genes. In other words, our analysis suggests that less than 2% of the cellular genes are not affected by EBV.

This view is supported by our time-resolved principal component analysis (Fig. 3A, D and E). It documents a well-defined status of the quiescent naïve B cells, which immediately changes upon EBV infection as a direct response to virus-mediated activating signals. Non-infected B cells express a characteristic set of genes that probably make up the signature of these quiescent naïve B-lymphocytes in vivo. This signature may support the cells in G0 and minimize their demand for energy, but upon infection many cellular genes show specific, time-regulated changes in their expression. Our bioinformatic work flow identified six distinct clusters (Fig. 4), which group them according to their expression pattern during infection and confirm our initial observations regarding cellular phenotypes and immunometabolism of the infected cells (Fig. 1).

The UpSet tool made it possible to identify and visualize the number of genes contained in unique as well as intersecting sets of differentially expressed genes between different days p.i.. Using this tool (Fig. 2C), we identified the most dynamic changes in host gene expression to occur very early after infection within 24 hours followed by considerable additional alterations within the next 48 hours, whereas later adjustments of the cells’ transcriptomes were relatively minor. Non-infected cells and cells infected for 24 hours formed a unique set of differentially expressed genes that does not intersect with any other set later during infection (Fig. 2B, C). Overall, our RNA-seq data analyses revealed that the most dynamic adjustments of gene expression occurred within the first four days after infection. When the infection progresses further EBV might need only to fine-tune the expression profile of the host cell to adjust the cell for the ensuing latent phase of infection.

### Viral gene expression precedes the time-controlled alterations in the cellular transcriptome

Scattered reports in the literature describe single viral genes and their early expression in B-lymphocytes (Price and Luftig, 2014). The results from our study mostly confirm the known or presumed kinetics of *in vitro* EBV gene expression in the pre-latent phase (Fig. 5). We identified the immediate and maximal induction of EBNA2, EBNA-LP, and BHRF1 on day 1 p.i., followed by a considerable reduction of their expression levels later. The expression of the EBNA3s peaks at day two p.i., whereas viral genes including LMP1 and LMP2s displayed a gradual increase in expression starting from day three p.i.. We also noticed that several viral genes defined as lytic genes are moderately expressed immediately after infection as expected (Kalla and Hammerschmidt, 2012), but become rapidly downregulated starting already on the second day p.i. (Fig. 5A; Supplementary Fig. S11A). Our observations support the asynchronous transcription of almost all viral genes immediately after the nuclear entry of viral DNA. Their rapid downregulation is probably mediated by early chromatin functions that shield the epigenetically naïve viral DNA from further promiscuous transcription. The strong and timely epigenetic repression of most viral genes (which spares EBV’s latent genes) is likely needed to establish and maintain latency (Woellmer et al., 2012; Woellmer and Hammerschmidt, 2013).

The profile of viral gene expression strongly correlates with the most prominent changes in the host B cell but highlights EBNA2 as the main driver together with EBNA-LP and BHRF1. EBNA2’s immediate and high expression induces Myc that presumably governs most subsequent alterations in the host B cell. This hypothesis is supported by an accompanying paper (Pich et al., 2019). In a systematic genetic approach with individual viral genes we identify EBNA2 to be the only viral gene that is indispensable for B cell activation, cell cycle entry and proliferation. We also noticed that not all naïve B-lymphocytes uniformly respond to EBV infection suggesting that EBV might specifically target non-identified B cell subpopulations or is incapable of activating all cells because viral or cellular gene expression is inconsistent. These open issues can only be addressed by single cell experiments (e.g. single cell RNA-seq), which will likely unveil how EBV reprograms individual target cells.

## Materials and Methods

Detailed descriptions of the materials and methods are provided in Supplementary Information of this paper and include the following:

Eukaryotic cell lines

Virus supernatants from the HEK293 2089 cell line

Collection and quantitation of virus supernatant from the B95-8 cell line

Preparation of B cells from adenoid tissue

Purification of B cells on MACS columns

Primary B cells infection with EBV

TMRE and Annexin V detection

Glucose uptake using 2-NBDG

FACS sorting for naïve B cells

Analysis of the cells’ diameter

Cell cycle analysis

Calculation of the cell division index (DI)

Preparation of cell lysates

Sample collection for time-course RNA-seq experiments

RNA isolation

Library preparation and sequencing

Bioinformatic analysis including transcript quantification by Salmon; quality control and data normalization; identification of differentially expressed genes; gene clustering; GO analysis; heatmaps of log10 transformed expression data; principal component analysis of phenotypic data; PCA with viral genes; clustering of viral genes; and data availability.

## Supplementary Information

Supplementary Information encompasses the section with details of Materials and Methods and eleven Supplementary Figures including their figure legends.

## Acknowledgements and Funding

We thank Christine Göbel, Munich, for her precious experimental support and expertise and Hannah Busen, Munich, for her advice and support during the initial phase of bioinformatic analyses. This work was financially supported by grants of the Deutsche Forschungsgemeinschaft [grant numbers SFB1064/TP A13, SFB-TR36/TP A04] to W.H., Deutsche Krebshilfe [grant number 70112875] to W.H., National Cancer Institute [grant number CA70723] to W.H‥

## Authors contributions

P. M.-G. designed and performed experiments, analyzed data, and wrote the paper. P.M.-G., A. B., T.S., R.F., and A.S. analyzed NGS data and performed the bioinformatic analyses. R.F. established the webpage. D.P. and W.H. performed experiments. W.H. conceived the project. W.H. and A.S. designed experiments and the bioinformatic workflow and wrote the paper.

