## Supplemental Text and Figures S1-S11 for "Epstein-Barr virus reprograms human B-lymphocytes immediately in the pre-latent phase of infection"

### Supplementary Information

Paulina Mrozek-Gorska[1], Alexander Buschle[1], Dagmar Pich[1], Thomas Schwarzmayr[2], Ron Fechtner[3], Antonio Scialdone[3,4,5*], and Wolfgang Hammerschmidt[1*]

[1]Research Unit Gene Vectors, Helmholtz Zentrum München, German Research Center for Environmental Health and German Center for Infection Research (DZIF), Partner site Munich
Marchioninistr. 25
D-81377 Munich, Germany

[2]Institute of Human Genetics, Helmholtz Zentrum München, German Research Center for Environmental Health
Ingolstaedter Landstr. 1
D-85764 Neuherberg, Germany

[3]Institute of Computational Biology, Helmholtz Zentrum München, German Research Center for Environmental Health
Ingolstaedter Landstr. 1
D-85764 Neuherberg, Germany

[4]Institute of Epigenetics and Stem Cells, Helmholtz Zentrum München, German Research Center for Environmental Health
Marchioninistr. 25
D-81377 Munich, Germany

[5]Institute of Functional Epigenetics, Helmholtz Zentrum München, German Research Center for Environmental Health
Ingolstaedter Landstr. 1
D-85764 Neuherberg, Germany

*Corresponding authors

### Materials and Methods

#### Eukaryotic cell lines

B95-8 (Miller et al., 1972), Raji (Pulvertaft, 1964), Elijah cells (Rowe et al., 1985), and LCLs (derived from EBV transformed B cells) were cultivated in RPMI-1640 medium supplemented with 8 % FCS, 100 µg/ml streptomycin, 100 U/ml penicillin, 1 mM sodium pyruvate, 100 nM sodium selenite, and 0.43 % α-thioglycerols at 37°C and 5 % $CO_2$. HEK293 2089 cells (Delecluse et al., 1998) were cultured in the same medium with 100 µg/ml hygromycin.

#### Virus supernatants from the HEK293 2089 cell line

HEK293 2089 in 13 cm dishes at around 80 % confluency were used for transient transfection with two plasmids, p509 and p2670, to induce EBV's lytic cycle. p509 expresses the BZLF1 gene under the control of the CMV promoter(Hammerschmidt and Sugden, 1988), which triggers lytic phase reactivation. p2670 contains the BALF4 gene under the CMV promoter to increase the virus titer (Neuhierl et al., 2002). 6 µg p509 and 6 µg p2670 in RPMI medium were mixed with PEI Max (linear polyethyleneimine hydrochloride, molecular weight 40,000, Polysciences Ltd, order number 24765-1) at a 1:6 w/w ratio. After 15 min incubation, the mix was dropped onto the HEK293 2089 cells prepared with fresh medium without selection and the cells were incubated for three additional days. The supernatants from the transfected cells were collected and centrifuged for 8 min at 300 g and at 1200 g for 8 min. The virus stocks were stored at 4°C and used to determine the virus concentration with Raji cells as described in detail (Steinbrück et al., 2015).

#### Collection and quantitation of virus supernatant from the B95-8 cell line

B95-8 cells were cultivated at high density to promote the spontaneous reactivation of EBV's lytic phase and virus release. Virus stocks were prepared as described above and virus titers were estimated using Elijah cells. The cells were incubated with different amounts of virus supernatant at 4°C for 3 hours and later stained with an Alexa 647 coupled antibody directed against the viral glycoprotein gp350. The fractions of gp350-positive cells with B95-8 virus stock were compared with cells incubated with a calibrated reference of 2089 virus by FACS.

#### Preparation of B cells from adenoid tissue

The adenoid biopsies were rinsed with PBS, transferred to a sterile petri dish, and mechanically disintegrated with two sterile scalpels and PBS. The cell suspension was filtered through a 100 µm mesh cell strainer. This procedure was repeated several times to recover a maximal number of single cells. The volume of the collected cells was increased to 50 ml with PBS, the cells were mixed and sedimented at 300 g for 10 min. The cell pellet was resuspended in 30 ml PBS and supplemented with 0.5 ml defibrinated sheep blood for T cells rosetting. 15 ml Ficoll-Hypaque was added underneath the cell suspension to obtain two clearly separated phases. The samples were centrifuged at 500 g for 30 min and the cells were carefully collected from the turbid interphase and transferred to a new 50 ml tube, which was filled with PBS. The cells were washed with PBS three times, using decreasing centrifugation parameters (450, 400 and 300 g for 10 min). Finally, the cell pellet was resuspended in fresh, pre-warmed medium. The cells were counted and immediately put to use.

**Purification of B cells on MACS columns**

All the steps were performed on ice and with pre-cooled solutions. Cells obtained from adenoid biopsies were centrifuged (300 g for 10 min) and washed once with ice-cold FACS staining buffer (PBS, 0.5 % BSA, 1 mM EDTA). Untouched B cells from human PBMCs were isolated according to the manufacturer's protocol using the B Cell Isolation Kit II (Milteny Biotec). The collected cell fractions were stained with an appropriate antibody marker (e.g. CD19 for B cells) to control the purity of the B cell purification.

**Primary B cells infection with EBV**

Infection of primary B cells was performed using wildtype EBV (2089 or B95-8) with a multiplicity of infection (MOI) 0.1 overnight. The next day the cells were centrifuged (300g at RT for 10 min) to remove unbound virus and were resuspended in fresh medium.

**TMRE and Annexin V detection**

For the detection of these parameters, cells from PBMC were purified using the B Cell Isolation Kit II (Milteny Biotec). Uninfected or infected B cells ($5x10^5$) were resuspended in 1 ml fresh pre-warmed medium and transferred to polypropylene FACS tubes. TMRE (100 nM final concentration) was added and the samples were incubated protected from light at 37°C for 30 min. The cells were washed two times with FACS buffer and resuspended in 500 µl Annexin V staining buffer together with APC-coupled Annexin V in the dark for 10 min. Daily measurements with a FACS Fortessa instrument were performed for the first eight days after EBV infection (Fig. 1A).

**Glucose uptake using 2-NBDG**

The experiments were performed with B cells from PBMC purified with the B Cell Isolation Kit II (Milteny Biotec). Non-infected cells and cells infected with wt EBV (B95-8) were collected on a daily basis as indicated (Fig. 1). About $5x10^5$ B cells were seeded into a 24-well plate and resuspended in 500 µl of glucose-free medium supplemented with 1 % FCS. The cells were incubated with the glucose analogue 2-NBDG (2-[N-(7-nitrobenz-2-oxa-1,3-diazol-4-yl) amino]-2-deoxy-D-glucose) (100 µM) for one hour or were left untreated (0 µM). Eventually, the cells were washed two times, resuspended in 300 µl ice-cold PBS and measured on a FACS Fortessa instrument using the FITC channel (Fig. 1B).

**FACS sorting for naïve B cells**

Primary B cells were sorted using a FACS Aria IIIu device to isolate naïve and resting human B-lymphocytes from adenoid biopsies. Briefly, after the Ficoll gradient the cells were washed once, resuspended in FACS staining buffer to a final concentration of $1x10^8$cells/ml and stained using fluorophore-coupled antibodies against CD38 (eBioscience, #25-0389-42) and IgD (BD Pharmingen, (#555778) for 60 min in the dark. The samples were washed once with ice-cold FACS staining buffer, resuspended in the same buffer, and filtered through a 35 µm mesh cell strainer to obtain single cell suspensions. Sorting was performed with a 70 µm nozzle, a velocity of about 8,000 events/second and a sorting mask of "4-way purity". The gating criteria included: (i) living cells, (ii) single cells, and (iii) CD38-negative and IgD-positive cells.

Prior to the subsequent analyses, viable, infected B cells were physically sorted as indicated in Fig. 1C using the 100 µm nozzle and a velocity of about 5,000 events/second with the sorting mask of "4-way purity".

**Analysis of the cells' diameter**

The diameter of the cells was recorded using uninfected cells and B cells infected with wt EBV at different days post infection as indicated in Supplementary Figure S3A with the Tali Image-Based Cytometer (Invitrogen). The obtained images were captured using the Quick Count program and default settings after recalibration with Tali Calibration Beads and further analyzed with the Image J software. Four images with about 200 cells each were analyzed at each time point. Defined objects with a circularity >0.75 and an aspect ratio <1.3 were identified as cells. The radius of each identified cell was determined according to the calibration beads with 1.9 mm radius on average, and the cells' diameter and cell volumes were calculated.

**Cell cycle analysis**

The sorted naïve B cells were infected with wt EBV (2089) at MOI 0.1. The next day cells were washed and around $5x10^5$ cells were seeded per time point. Prior to cell harvest, the cells were always incubated with 3.1 µl BrdU dilution (1:32) per 100 µl of cells and incubated at 37°C for 1 hour in the dark. Afterwards, the cells were harvested and washed with 1 ml of FACS staining buffer. The cell pellet was resuspended in 100 µl of BD Cytofix/Cytoperm buffer followed by incubation for 15-30 min on ice. Next, the cells were washed with 1 ml FACS staining buffer, then frozen in freezing medium (90% FCS, 10% DMSO) and stored at -80°C for further analysis. After collecting all samples, the cells were thawed, re-fixated with 100 µl BD Cytofix/Cytoperm buffer for 5 min on ice, washed with 1 ml BD Perm/Wash buffer, and subsequently resuspended in 70 µl PBS and 30 µl of DNase. The mix was incubated at 37°C for 1 h, the cells were washed with 1 ml BD Perm/Wash buffer, resuspended in 50 µl BD Perm/Wash buffer and stained with 1 µl anti-BrdU-APC coupled antibody for 20 min at RT. After staining, the cells were washed with BD Perm/Wash buffer, resuspended in 20 µl 7-AAD solution and 500 µl FACS staining buffer. After 10 min the samples were measured with a FACS Fortessa instrument to determine the cell cycle distribution of the cells (Fig. 1D).

**Calculation of the cell division index (DI)**

Sorted naïve B cells were resuspended in warm PBS at a concentration $10^6$ cells/ml and mixed with diluted CellTrace violet (ThermoFischer Scientific, 1 µM final concentration). The cells were incubated in a water bath at 37°C and protected from light for 20 min. Afterwards, 50 ml of complete, pre-warmed culture medium was added to the cells to quench the unbound dye followed by 5 min incubation at 37°C. Finally, the cells were spun down at 300 g for 10 min and resuspended in fresh, pre-warmed medium. Prepared cells were infected with two wild-type EBV strains (2089 and 6008) at MOI 0.1 and incubated overnight. On the next day, the cells were spun down, resuspended in fresh medium and aliquoted for daily measurements depending on the initial number of cells. Measurements were performed daily on a FACS Fortessa instrument to determine the dilution of CellTrace violet dye as an indicator of cell division. The division index DI documents the average number of cell divisions that cells in the starting population have undergone (Supplementary Fig. S3B). The index was calculated using the FlowJo Proliferation Tool software.

**Preparation of cell lysates**

Preparation of whole cell extracts was performed with exactly $1x10^6$ uninfected cells or with cells collected at different time points after EBV infection. The cells were washed once with PBS (6 min at 300 g and 4°C) and lysed in a volume of 200 µl RIPA buffer for 30 minutes on ice. The lysates were sonicated with a Bioruptor sonicator (30 sec on/30 sec off, 'high' setting, 4 x 5 min, on ice). Afterwards, the samples were centrifuged to remove cellular debris (20 min at maximal speed at 4°C) and the supernatants were transferred to new tubes. The protein concentrations of the lysates were determined using the Bradford's protein assay (EMD Millipore) (Fig. 1E). The absorbance was measured at 595 nm in a photometer and protein concentrations were determined using a BSA standard curve.

**Sample collection for time-course RNA-seq experiments**

B cells were purified from adenoid biopsies and sorted to enrich for naïve B cells. The "Day 0" samples with $1x10^6$ uninfected naïve B cells were washed two times with PBS and resuspended in 1 ml Trizol reagent. The samples were snap frozen in liquid nitrogen and stored for further analysis at -80°C. The remaining cells were bulk infected (MOI 0.1) with wt EBV (2089). On the next day, the cells were spun down to remove unbound viral particles and resuspended in fresh medium supplemented with Ciprobay (1:200, Bayer) and Cyclosporine A (1 µg/ml, Sigma). Cells were aliquoted for the harvest on different days after infection. At the indicated time points (Fig. 2) the cells were collected and resuspended in FACS staining buffer. Living cells were sorted according to the established gating strategy to obtain one million cells. The cells were washed two times with PBS, resuspended in Trizol reagent and snap frozen in liquid nitrogen. Frozen cells were further used for RNA extraction.

**RNA isolation**

Total RNA was extracted from $1x10^6$ sorted and frozen samples (day zero to five, day eight, and day 14p.i.). Briefly, the samples in Trizol were thawed on ice and mixed with 200 µl chloroform, vortexed and centrifuged with maximal speed for 20 min at 4°C. The aqueous phase was carefully transferred to a new sterile RNase-free tube and combined with an equal volume of RNA-free absolute ethanol. Samples were then loaded onto RNeasy columns and total RNA extraction was performed using the RNAeasy Kit (Qiagen) including DNase treatment. Total RNA quality and concentration were measured by capillary electrophoresis using the Shimadzu MultiNA Electrophoresis System for DNA/RNA analysis (Fig. 1F).

**Library preparation and sequencing**

Different amounts of RNA were used (30 ng from Day0, 50 ng from Day1, 100 ng from Day2, and 200ng from samples collected at Day3-5, 8, and 14) to generate the sequencing libraries. The first-strand cDNA was synthesized using oligo(dT) primer. cDNA samples were fragmented and Illumina TruSeq sequencing adapters were ligated to the 5´ and 3´ends of the cDNA fragments. Barcoded cDNA samples were finally amplified by PCR using a proof-reading polymerase. The primers used for PCR amplification were designed for TruSeq Dual-Index sequencing according to the instructions by Illumina. Aliquots of size-selected cDNAs (200-600 bp) were measured by capillary electrophoresis to confirm the correct size and to determine the concentration of each sample. All prepared libraries were sequenced together (paired-end, 100 bases) using two entire flow cells of the Illumina HiSeq4000 platform (Institute of Human Genetics, Helmholtz Zentrum München).

**Bioinformatic analysis**

**Transcript quantification by Salmon**

Transcripts were quantified by running Salmon version v0.9.1 (Patro et al., 2017) in the quasi-mapping-based mode. First, a transcriptome index was created for the set of human reference transcripts (GRCh37.p13) and a set of (manually assembled) 2089 EBV transcripts. Then, the quantification step was carried out with the "quant" function, correcting for the sequence-specific biases ("--seqBias" flag) and the fragment-level GC biases ("--gcBias" flag). Finally, the transcript level abundances were aggregated to gene level counts.

**Quality control and data normalization**

As a first assessment of sample quality, the library size, the number of detected genes, and the ratio between the reads mapped to viral genes and those mapped to endogenous genes were computed. The quality control identified very consistent library size and number of detected genes between samples but for sample 8_1 (Supplementary Fig. S4). An additional PCA analysis confirmed the bias in the sample 8_1, which was excluded from further analysis.

All the samples that passed the quality control were normalized for sequencing depth using size factors (Anders and Huber, 2010). After normalization, a principal component analysis (PCA) was performed on the log-transformed expression matrix of all genes (Fig. 2A).

**Identification of differentially expressed genes**

The DESeq2 (Love et al., 2014) R package (version 1.16.1) was used to identify differentially expressed genes between pairs of samples from different days post infection. Before running DESeq2, genes with an average expression lower than 50 normalized counts were excluded. Next, all genes that were significantly differentially expressed (DE) at a false discovery rate <0.1 and had an estimated fold-change of >2 or <0.5 in at least one pair-wise comparison were selected. This resulted in a set of 11,178 DE genes, which were then clustered (see below).

Among all analyzed genes with read counts ≥50, only 241 genes were not significantly differentially expressed between any of the time points after infection, suggesting that these genes were not regulated during infection. This number corresponds to 1.85 % of all genes tested (n=13,007).

MA plots visualizing the log fold change of gene expression versus mean normalized counts (Supplementary Fig. S7) were obtained during DESeq2 analysis. The intersections of DE genes between any given pair of time points was visualized by UpSet plots (Lex et al., 2014)(UpSetR R package, version 1.3.3).

**Gene clustering**

The list of 11,178 DE genes identified above were clustered according to their expression patterns along the infection time course. We calculated a distance matrix between genes as:$\sqrt{(1-\rho)/2}$, where $\rho$ is the Spearman's correlation coefficient between pairs of genes across all samples (van Dongen and Enright, 2012). Hierarchical clustering was performed on this distance matrix ("hclust" function in R, with the "average" aggregation method), followed by the dynamic hybrid cut algorithm to estimate the number and identity of clusters (dynamicTreeCut package v1.63-1, with minimum cluster size of 500 and "deepSplit" parameter equal to 0, as suggested by our robustness analysis, see below). The analysis resulted in six clusters with specific expression patterns of cellular genes along the time-course of EBV infection (Fig. 4).

We assessed the robustness of the clusters obtained from the dynamic hybrid cut algorithm at all possible values of the "deepSplit" parameter. To do so, we computed the values of the Pearson Gamma and the average Silhouette width of the different clustering; both these parameters suggested that the best clustering was obtained with deepSplit=0, as indicated by the location of their maximum (Supplementary Fig. S8B,C).

We also checked how robust the clusters were to gene sub-setting. More specifically, for each value of "deepSplit", we clustered the dataset 100 times after randomly removing 10% of genes and computed the values of the Pearson Gamma and the average Silhouette width (Supplementary Fig. S8D,E). Again, the most robust clusters were found with deepSplit=0. The statistics of Pearson Gamma and Silhouette width were calculated with the "clust.stats" function from the "fpc" R package (version 2.1-11.1).

The six gene clusters were visualized with a t-SNE representation ("RtSNE" R package version 0.13) of the distance matrix with perplexity set to 50 (Supplementary Fig. S8A).

**GO analysis**

Each group of genes assigned to one of the six clusters was analyzed with the gene ontology (GO) online tool (GOrilla) (Eden et al., 2009)to characterize specific biological processes that are enriched in each cluster with respect to the list of all DE genes.

To increase the specificity of the enriched GO term we set the p-value threshold to $\leq 10^{-3}$ and harmonized the redundancy in identified GO terms by additional filtering with the "reduce and visualize Gene Ontology" (REViGO) online tool with the following settings: medium similarity, database for Homo sapiens, and SimRel for calculating the semantic similarity of the GO terms (Supek et al., 2011). R scripts created by the REViGO tool were employed to create tree plots from the enriched GO terms for each of the six clusters (Supplementary Fig S9).

**Heatmaps of log10 transformed expression data**

Heatmaps were generated using "pheatmap" R package version 1.0.10 with default settings on log10 transformed expression data of the top 100 cellular and viral genes (Supplementary Fig. S6) or only viral genes (Fig. 5B) contributing to the first two principal components.

**Principal Component Analysis of metabolic and phenotypic data**

The Principal Component Analysis (PCA) matrix was calculated based on the normalized means of the results derived from seven different read-outs: TMRE binding, 2-NBDG uptake, cell diameter, cell division index, S-phase fraction of cells, protein content, and total RNA content. The results of TMRE uptake were expressed as the ratio of the TMRE_high positive cell population versus all TMRE positive cells (TMRE_highplusTMRE_low). The fraction of cells that actively sequester 2-NBDG and the fraction of cells in S phase were calculated. The average from three to four biological replicates comprising cell diameter, division index, protein content, and total RNA content were normalized according to the maximal value identified in the individual experiments. After normalization, the matrix with all data from the seven different experiments was used to perform the PCA in R. The first two principal components (PC1 and PC2) were plotted (Fig. 1G) to identify the directions explaining most of the variance in the metabolic and phenotypic data.

**PCA with viral genes**

A principal component analysis (PCA) was performed in R on the matrix of log-transformed normalized read counts of viral genes, excluding the day 0 time point, where no viral genes were expressed (Supplementary Fig. S10A).

**Clustering of viral genes**

From the significantly DE genes identified by DESeq2, we selected the 27 DE viral genes and considered them for clustering. A distance matrix was created as explained above (see the previous paragraph “Gene clustering”). In the next step, hierarchical clustering (“hclust”) and dynamic hybrid cut algorithm (“dynamicTreeCut”) were used on the distance matrix (minimum cluster size equal to 5 and deepSplit=0) to estimate the identity and the number of clusters. We found three clusters that were named with Roman numerals. The expression values of viral genes were plotted as mean normalized expression versus time-points (days p.i.) after EBV infection (Supplementary Fig. S10B).

**Data availability**

Raw RNA-seq data are available at – URL information needs to be added here, eventually. Processed RNA-seq data can be downloaded from the following website, which also provides a user-friendly R Shiny App to visualize and to explore the data:

https://scialdonelab.shinyapps.io/EBV_B/

### Supplementary Figure legends

#### Supplementary Figure S1. Annexin V binding and mitochondrial activity during the first eight days after EBV infection.

The FACS plots show the rate of early apoptosis (Annexin V) and mitochondrial activity (TMRE) of the cells at early time points after EBV infection. The percentages of positive cells as measured by FACS are indicated.

Panel **A** shows established LCLs as a control, panel **B** displays uninfected primary B cells (Day0) or cells on different days p.i.. In the first column, the forward and sideward scatter of the different cell samples are compared. The next two columns analyze untreated cells (0 nM) and cells incubated with 100 nM TMRE for 30 min. Columns four to six show the results after back-gating of the cell populations indicating the fraction of cells that were Annexin V positive (red), TMRE_low positive (blue) or TMRE_high positive (green). TMRE incorporation revealed two clearly positive populations in uninfected B cells (Day0). The population of TMRE_high cells became dominant from day four p.i. indicating a high mitochondrial activity of stably infected cells. The figure shows the results of one representative experiment out of four biological replicates.

#### Supplementary Figure S2. Uptake of the glucose analogue (2-NBDG) during B cells infection with EBV.

FACS analysis of the glucose uptake of B cells at different days after EBV infection. Panel **A** shows an established LCL as a reference, the remaining **B** panels represent uninfected B cells and cells infected for different time periods as indicated. The first column compares forward and sideward scatter of cells incubated with 2-NBDG. The next three columns show the analysis of untreated cells (0 µM 2-NBDG) and cells incubated with 100 µM of 2-NBDG. During the first three days after viral infection the cells did not show significant changes in glucose uptake. Starting from day four p.i., the glucose uptake increased substantially to a fraction of more than 85 % cells with high glucose uptake. The figure depicts one experiment out of four performed with B cells from different donors.

#### Supplementary Figure S3. Cell size and dynamics of cell divisions during EBV infection.

**A.** The panel shows the changes of the cellular diameter during the first eight days after EBV infection. Mean and standard deviation were calculated from the images of about thousand cells at the specified time points p.i..

**B.** Sorted naïve B cells were treated with the intracellular dye CellTrace Violet (CTV) and subsequently infected with two wild type strains of EBV at MOI 0.1. On a daily basis, the calls were analyzed by FACS and based on the dilution of the CTV the Division Index (DI) was determined and plotted. The graph shows one representative example out of three experiments with different B cell donors.

**Supplementary Figure S4. Quality control of sequencing libraries.**

**A.** The sizes of the sequenced libraries prepared from the 24 different samples from three independent donors are depicted. The majority of the samples shows comparable library sizes except sample Day8_1, which was removed from the downstream analysis.

**B.** The bar plot depicts the numbers of detected genes with a threshold of >10 reads per million (RPM) in each sequenced sample. The numbers of detected genes were comparable.

**C.** The ratios between the number of reads allocated to viral genes and numbers of reads allocated to cellular genes at each day for three replicates are shown. Viral genes were absent in uninfected cells (Day0) in contrast to day one and two, when their relative abundance reached a maximum. From day three on, the abundance of EBV genes decreased to reach a stable level at day four p.i.. Once again, Sample Day8_1 appears to be an outlier because of its high ratio of viral versus cellular genes.

**Supplementary Figure S5. Schematic workflow of the bioinformatic analyses after RNA-seq.**

The scheme shows the individual steps of the bioinformatic workflow starting with the raw data analysis and transcript quantification, followed by data normalization and PCA analysis. Finally, the DE analysis together with an Upset intersection analysis and a gene clustering step concluded the analyses (details of the single steps of data analyses can be found in Materials and Methods).

**Supplementary Figure S6. Heatmaps of the top 100 genes that contribute to PC1 or PC2.**

We considered 50 genes with the most positive and 50 genes with the most negative loadings on the first two principal components (PC1 and PC2; relative to the PCA plot shown in Figure 2A). The log10 transformed mean expression values of the combined 100 genes (rows) were plotted in the heatmap at different days after EBV infection (columns). Panel **A** includes genes selected from PC1 and panel **B** displays genes contributing to PC2. A color gradient from yellow to red indicates expression values from the lowest to the highest, respectively.

**Supplementary Figure S7. MA plots of gene expression on consecutive days after EBV infection.**

The MA plots show the comparison of gene expression levels between consecutive days p.i. (Day0 vs Day1, Day1 vs Day2, etc.). The x-axes display the means of normalized counts and the y-axes depict the log10-fold changes between the time points indicated above the plot. Every dot represents a single gene and genes that passed the significance threshold (FDR) <0.1 are shown in red.

**Supplementary Figure S8. Clustering robustness analysis.**

**A.**t-SNE plot of all identified DE genes in the course of EBV infection. The colors indicate the six clusters estimated by the hierarchical clustering and the dynamic hybrid cut algorithm.

**B. and C.** The dynamic hybrid cut algorithm was used with all possible values of the "deepSplit" parameter with all DE genes. The values of the Pearson Gamma (PG) (B) and silhouette width (silwidth) (C) were calculated using the R function "cluster.stats" from the "fpc" package and plotted against the different values of the "deepSplit" parameters. Higher values of PG and silwidth correspond to a better clustering. Here, PG and silwidth achieve their maximum values at "deepSplit"=0 suggesting that the best clustering is obtained for such a value of "deepSplit".

**D. and E.** The same analysis as in panel B and C was performed with 100 randomly selected subsamples of genes. The dynamic hybrid cut was performed and clustering quality was assessed by computing the Pearson Gamma and silhouette width parameters. The x-axis of the plots depicts all used "deepSplit" values, whereas the y-axis displays values for PG (D) or silwidth (E). Also this analysis suggests that "deepSplit"=0 gives the most robust clusters.

**Supplementary Figure S9. Visualization of cluster-specific GO enrichments.**

The tree maps represent the GO enrichment analyses after reducing its redundancy using REViGO (http://revigo.irb.hr/). The plots contain rectangles of different sizes indicating the absolute -log10 of the calculated p-values of the shown GO terms. The color background of the tree maps relates to the color name of the clusters as indicated: the brown cluster (panel A), the green cluster (panel B), the yellow cluster (panel C), the blue cluster (panel D), the red cluster (panel E), and the turquoise cluster (panel F).

**Supplementary Figure S10. Viral gene analysis in the course of EBV infection.**

**A.** Principal Component Analysis (PCA) performed on viral genes. Each dot represents a biological replicate and different colors show different time points after EBV infection as indicated. The percentage of variance explained by the first two principal components is indicated in parentheses.

**B.** The dynamics of viral gene expression in each of the clusters is shown. The hierarchical clustering analysis (see Figure 3) determined three main clusters of viral genes (I, II, and III), which display a dynamic expression during the early phase of EBV infection. The plots illustrate the mean normalized expression levels (y-axis) versus days p.i. (x-axis). The number N of the genes in each cluster is indicated at the top of each plot and the insets contain the lists of the viral genes included in the depicted cluster.

**Supplementary Figure S11. Viral genes expression.**

The bar plots represent the expression levels (y-axis, normalized read counts) during the course of EBV infection (x-axis) of all the individual viral genes for cluster I (panel A), cluster II (panel B), and cluster III (panel C).

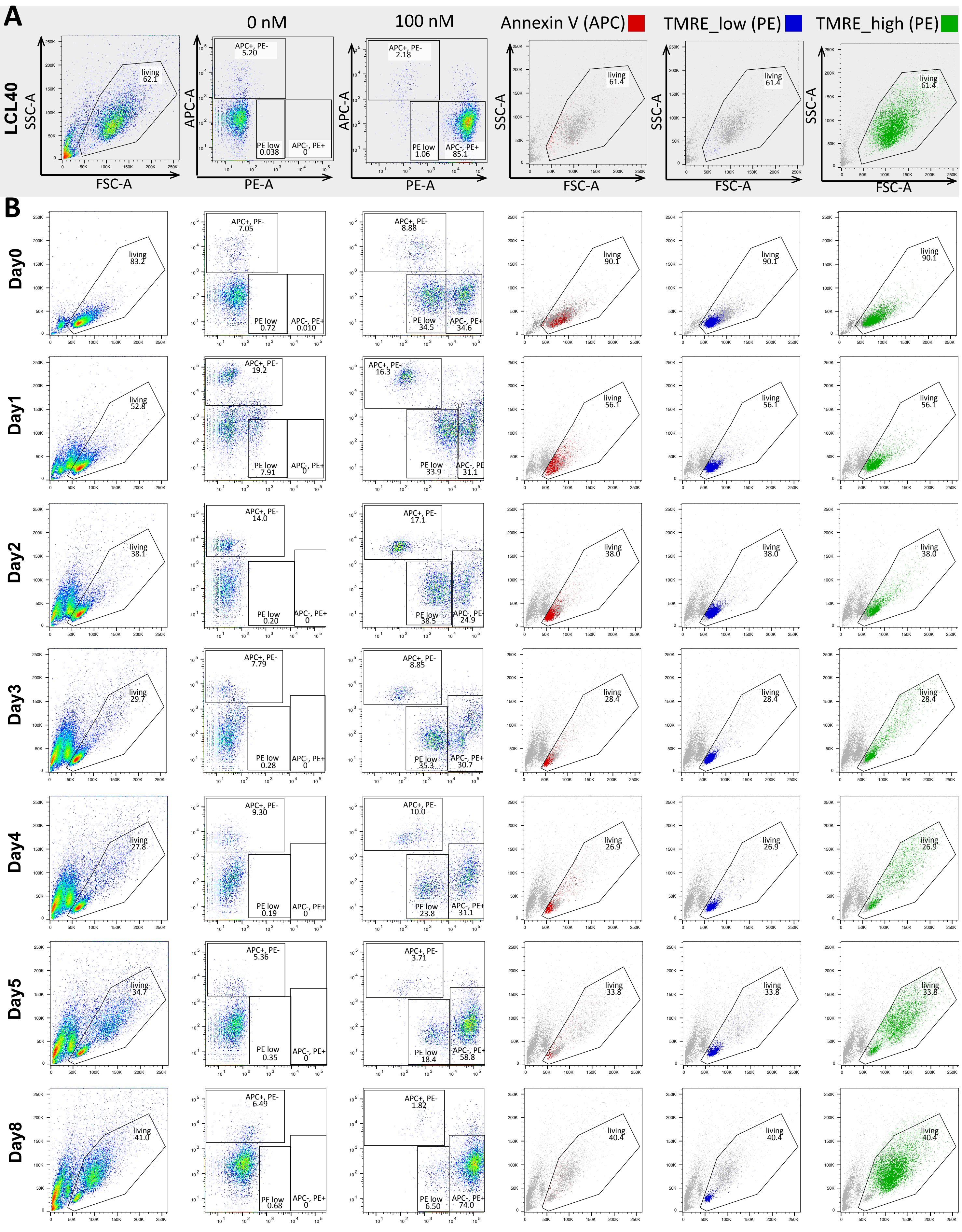
A
LCL40
0 nM
100 nM
Annexin V (APC)
TMRE_low (PE)
TMRE_high (PE)
SSC-A
FSC-A
APC-A
PE-A
living
APC+, PE-
PE low
APC-, PE+
B
Day0
Day1
Day2
Day3
Day4
Day5
Day8

Figure S1

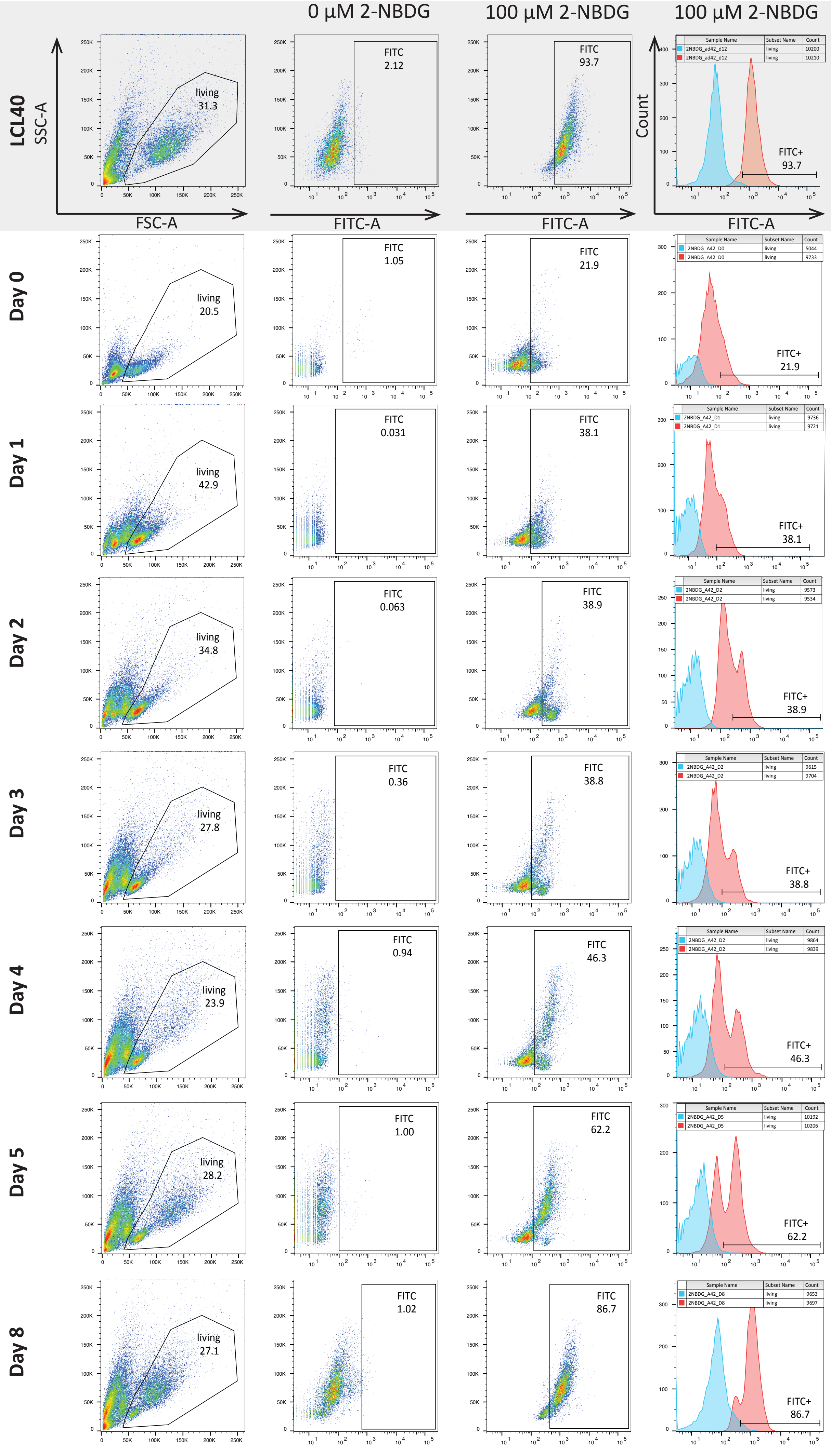
0 µM 2-NBDG
100 µM 2-NBDG
100 µM 2-NBDG
LCL40
SSC-A
FSC-A
FITC-A
Count
living 31.3
FITC 2.12
FITC 93.7
FITC+ 93.7
Day 0
living 20.5
FITC 1.05
FITC 21.9
FITC+ 21.9
Day 1
living 42.9
FITC 0.031
FITC 38.1
FITC+ 38.1
Day 2
living 34.8
FITC 0.063
FITC 38.9
FITC+ 38.9
Day 3
living 27.8
FITC 0.36
FITC 38.8
FITC+ 38.8
Day 4
living 23.9
FITC 0.94
FITC 46.3
FITC+ 46.3
Day 5
living 28.2
FITC 1.00
FITC 62.2
FITC+ 62.2
Day 8
living 27.1
FITC 1.02
FITC 86.7
FITC+ 86.7

Figure S2

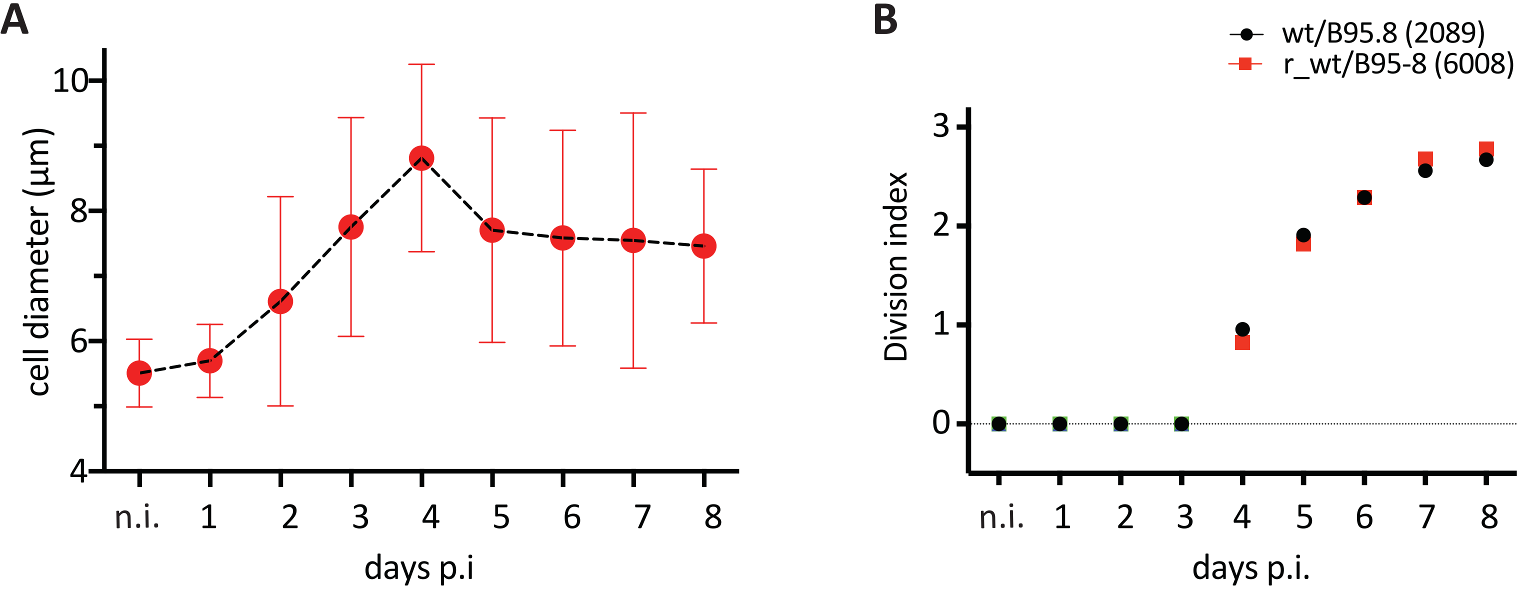
A
cell diameter (µm)
10
8
6
4
n.i.
1
2
3
4
5
6
7
8
days p.i
B
wt/B95.8 (2089)
r_wt/B95-8 (6008)
Division index
3
2
1
0
n.i.
1
2
3
4
5
6
7
8
days p.i.

Figure S3

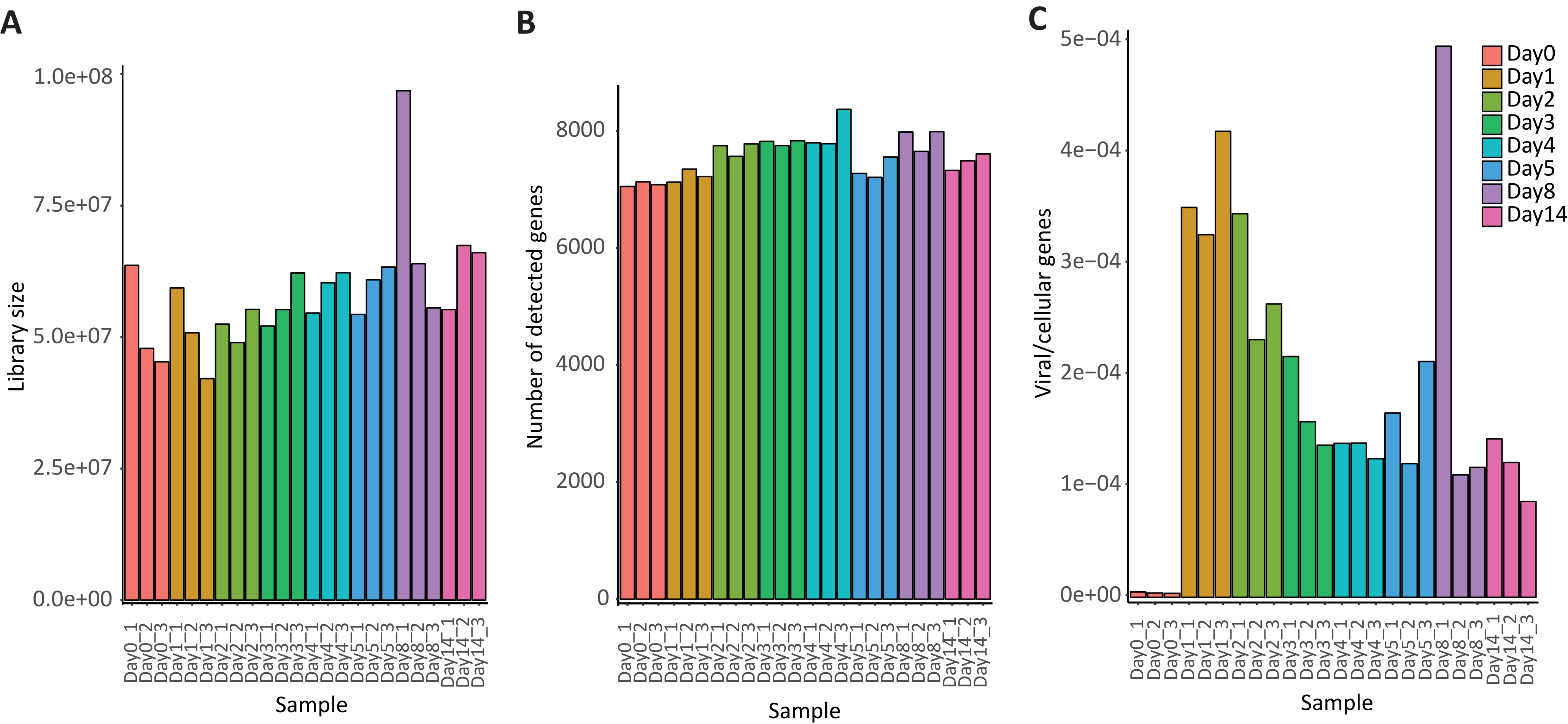
A
B
C
Library size
1.0e+08
7.5e+07
5.0e+07
2.5e+07
0.0e+00
Number of detected genes
8000
6000
4000
2000
0
Viral/cellular genes
5e−04
4e−04
3e−04
2e−04
1e−04
0e+00
Day0
Day1
Day2
Day3
Day4
Day5
Day8
Day14
Day0_1
Day0_2
Day0_3
Day1_1
Day1_2
Day1_3
Day2_1
Day2_2
Day2_3
Day3_1
Day3_2
Day3_3
Day4_1
Day4_2
Day4_3
Day5_1
Day5_2
Day5_3
Day8_1
Day8_2
Day8_3
Day14_1
Day14_2
Day14_3
Sample

Figure S4

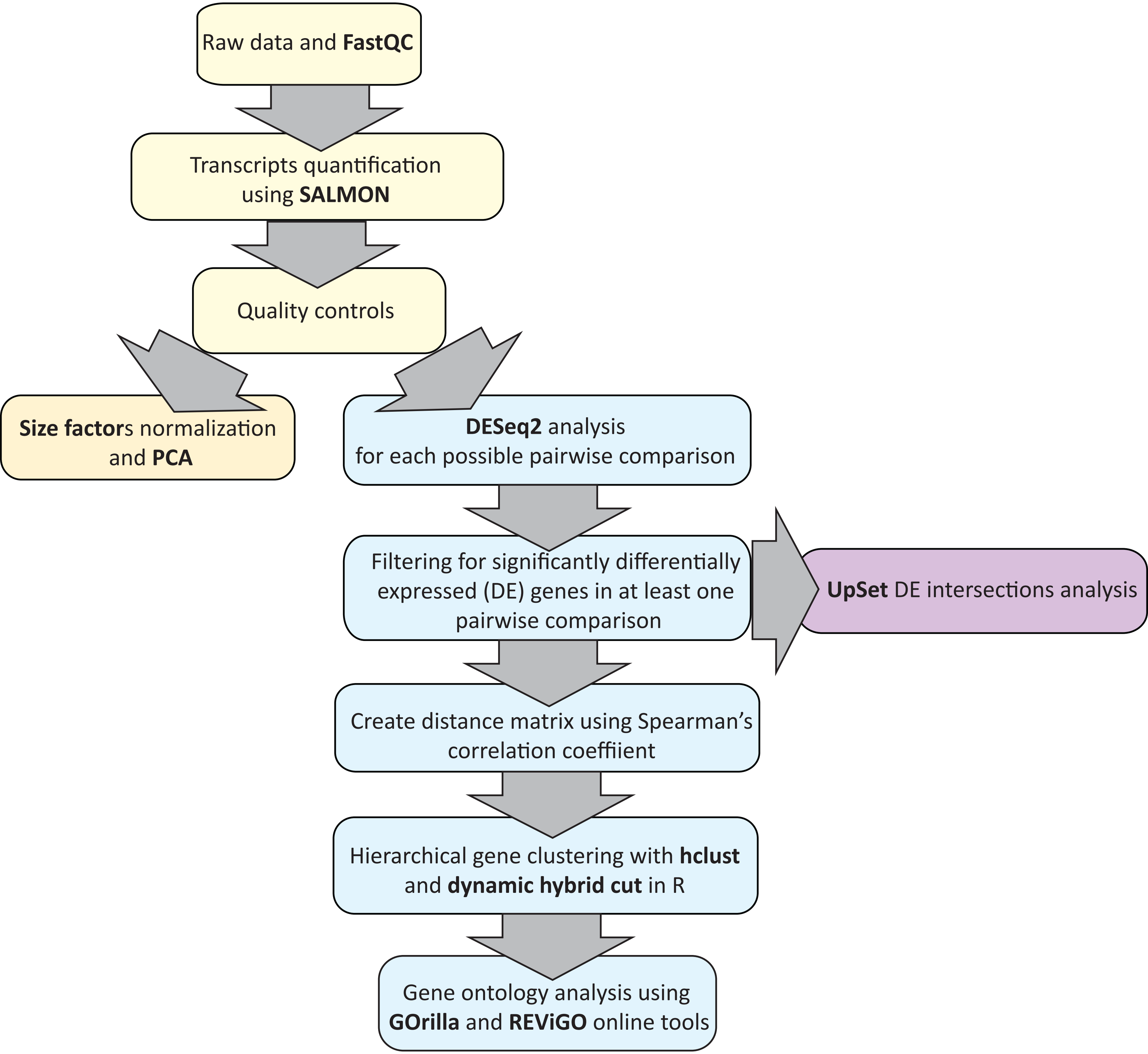
Raw data and FastQC
Transcripts quantification using SALMON
Quality controls
Size factors normalization and PCA
DESeq2 analysis for each possible pairwise comparison
Filtering for significantly differentially expressed (DE) genes in at least one pairwise comparison
UpSet DE intersections analysis
Create distance matrix using Spearman's correlation coeffiient
Hierarchical gene clustering with hclust and dynamic hybrid cut in R
Gene ontology analysis using GOrilla and REViGO online tools

Figure S5

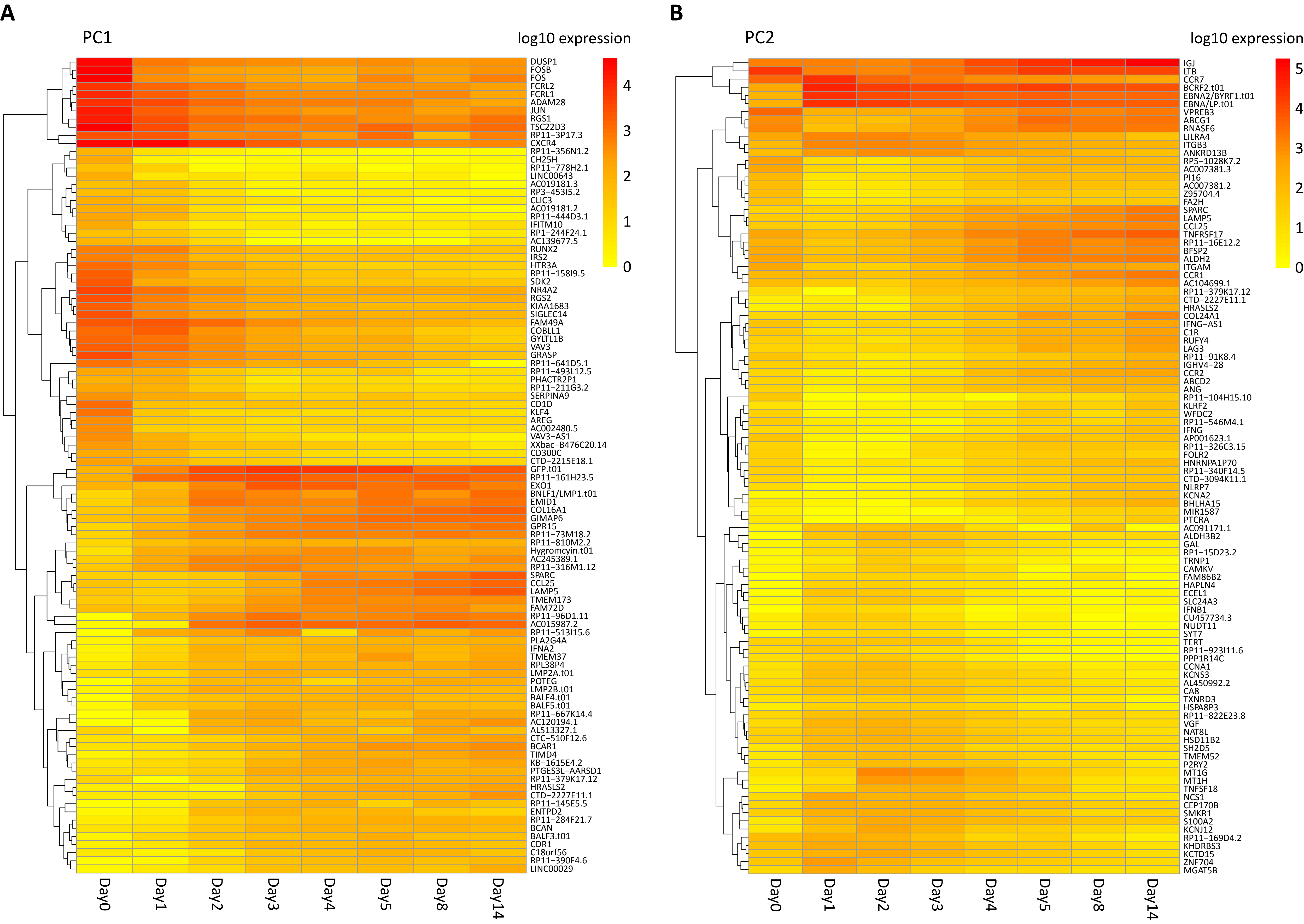
A
PC1
log10 expression
B
PC2
log10 expression
Day0
Day1
Day2
Day3
Day4
Day5
Day8
Day14

Figure S6

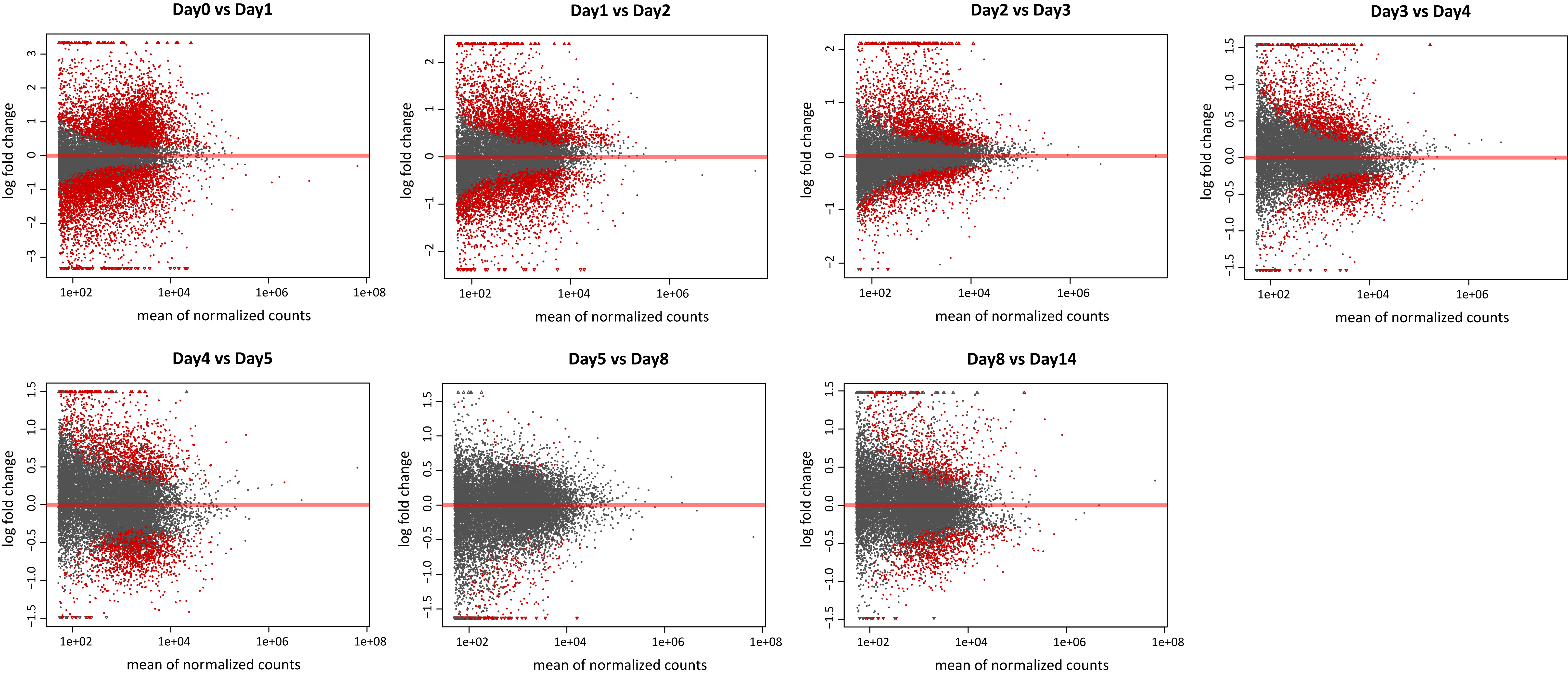
Day0 vs Day1
Day1 vs Day2
Day2 vs Day3
Day3 vs Day4
Day4 vs Day5
Day5 vs Day8
Day8 vs Day14
log fold change
mean of normalized counts
1e+02
1e+04
1e+06
1e+08

Figure S7

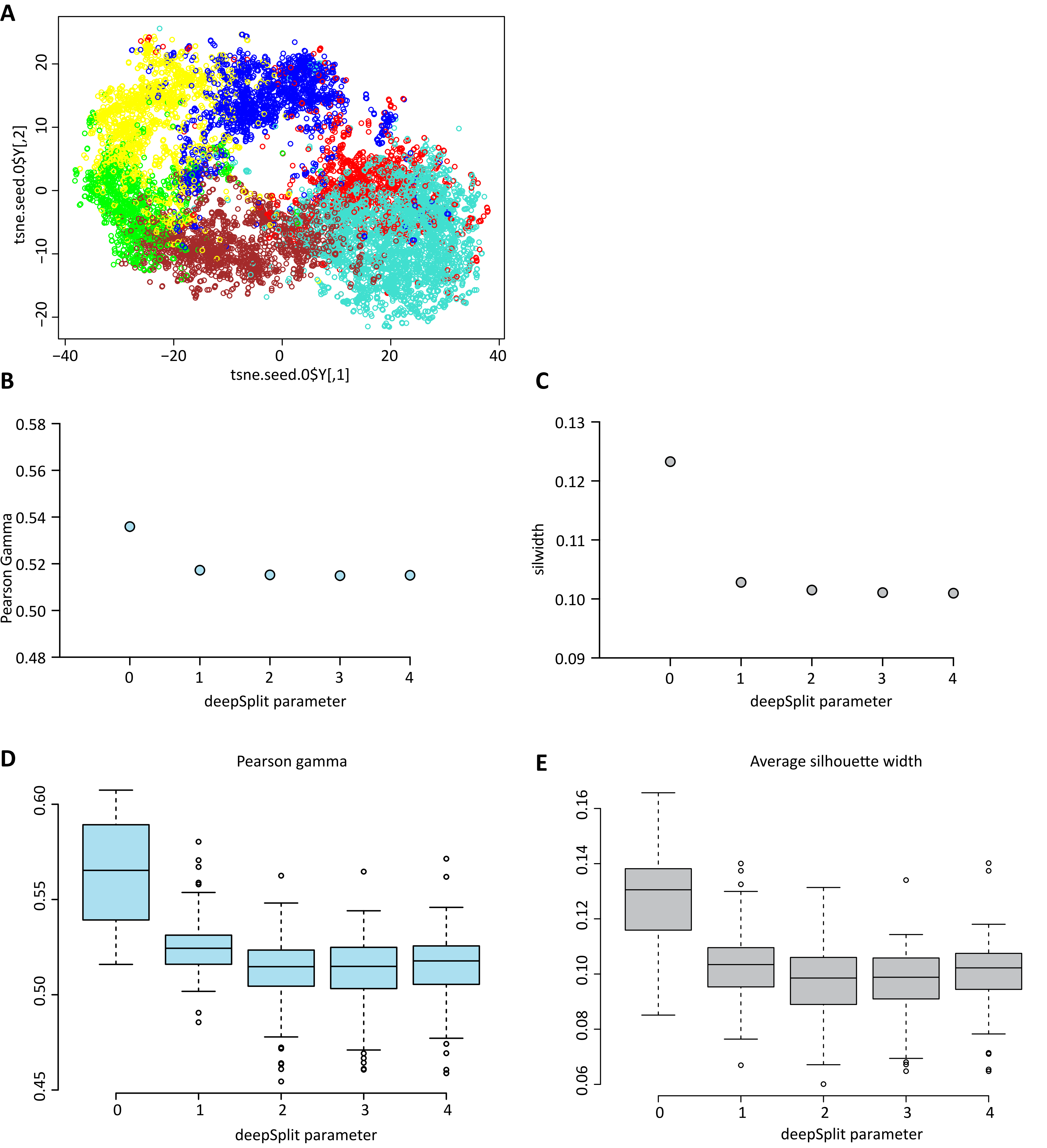
A
tsne.seed.0$Y[,2]
tsne.seed.0$Y[,1]
B
Pearson Gamma
deepSplit parameter
C
silwidth
deepSplit parameter
D
Pearson gamma
deepSplit parameter
E
Average silhouette width
deepSplit parameter

Figure S8

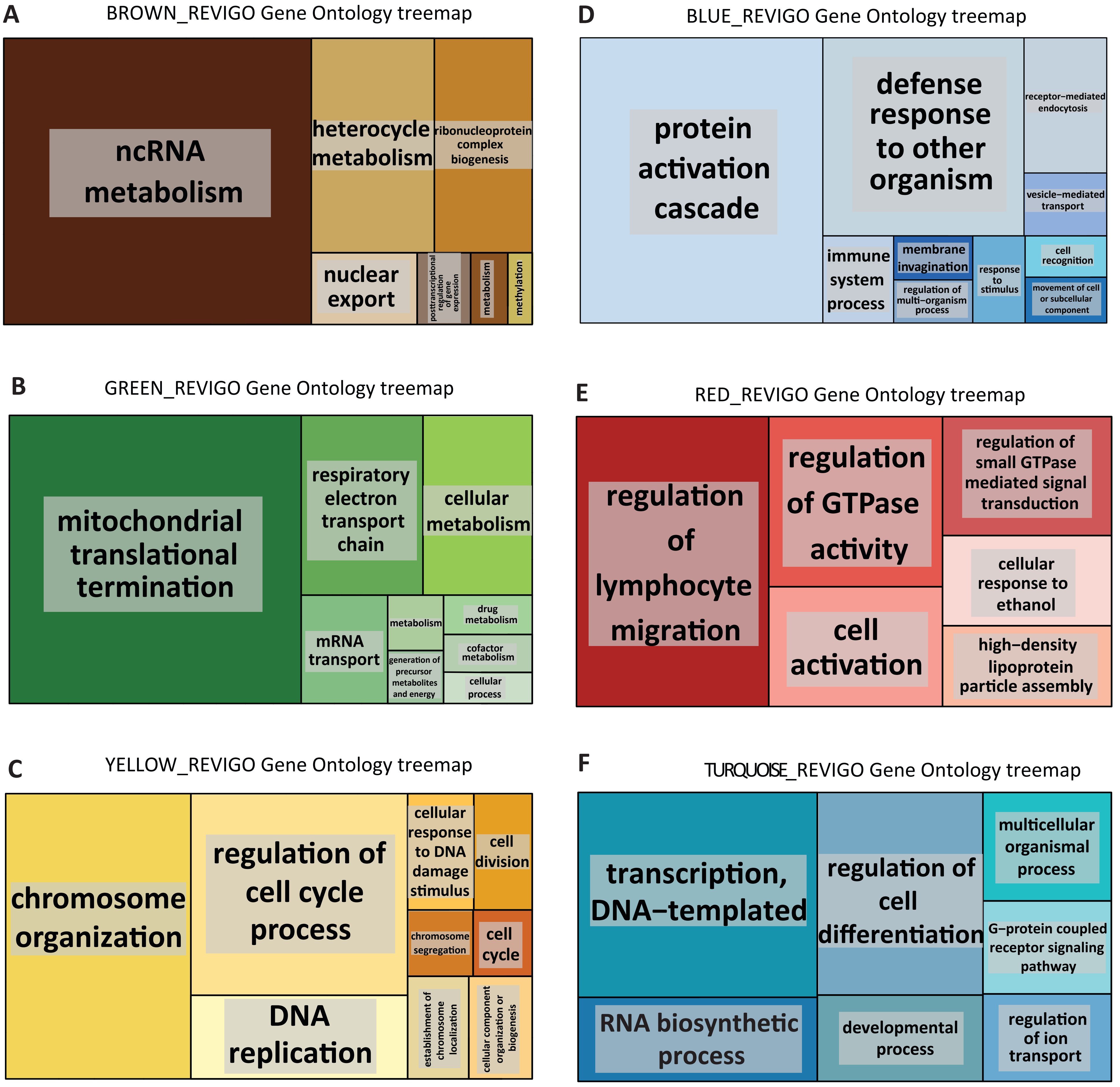
A
BROWN_REVIGO Gene Ontology treemap
ncRNA metabolism
heterocycle metabolism
ribonucleoprotein complex biogenesis
nuclear export
posttranscriptional regulation of gene expression
metabolism
methylation
B
GREEN_REVIGO Gene Ontology treemap
mitochondrial translational termination
respiratory electron transport chain
cellular metabolism
mRNA transport
metabolism
generation of precursor metabolites and energy
drug metabolism
cofactor metabolism
cellular process
C
YELLOW_REVIGO Gene Ontology treemap
chromosome organization
regulation of cell cycle process
DNA replication
cellular response to DNA damage stimulus
cell division
chromosome segregation
cell cycle
establishment of chromosome localization
cellular component organization or biogenesis
D
BLUE_REVIGO Gene Ontology treemap
protein activation cascade
defense response to other organism
receptor−mediated endocytosis
vesicle−mediated transport
immune system process
membrane invagination
regulation of multi−organism process
response to stimulus
cell recognition
movement of cell or subcellular component
E
RED_REVIGO Gene Ontology treemap
regulation of lymphocyte migration
regulation of GTPase activity
regulation of small GTPase mediated signal transduction
cellular response to ethanol
cell activation
high−density lipoprotein particle assembly
F
TURQUOISE_REVIGO Gene Ontology treemap
transcription, DNA−templated
regulation of cell differentiation
multicellular organismal process
G−protein coupled receptor signaling pathway
RNA biosynthetic process
developmental process
regulation of ion transport

Figure S9

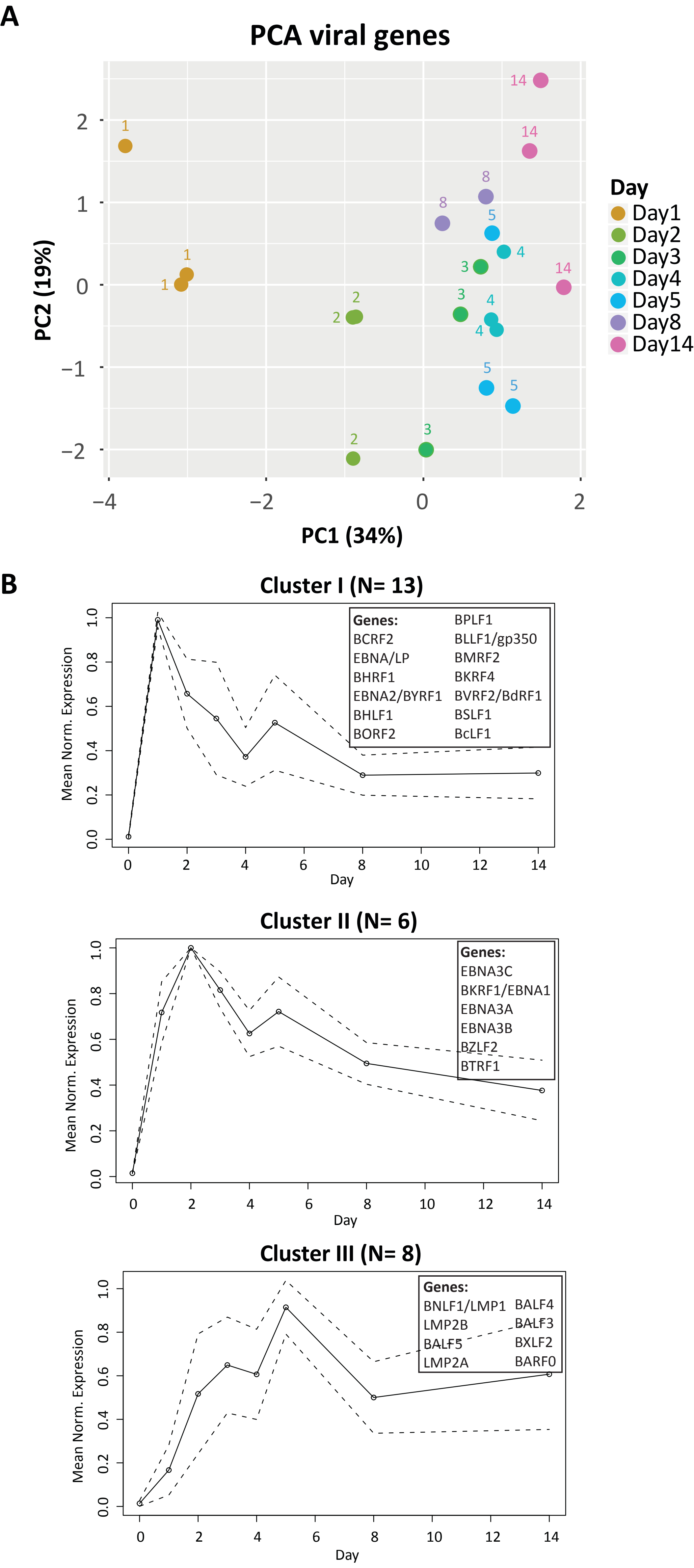
A
PCA viral genes
PC2 (19%)
PC1 (34%)
Day
Day1
Day2
Day3
Day4
Day5
Day8
Day14
B
Cluster I (N= 13)
Mean Norm. Expression
Day
Genes:
BCRF2
EBNA/LP
BHRF1
EBNA2/BYRF1
BHLF1
BORF2
BPLF1
BLLF1/gp350
BMRF2
BKRF4
BVRF2/BdRF1
BSLF1
BcLF1
Cluster II (N= 6)
Mean Norm. Expression
Day
Genes:
EBNA3C
BKRF1/EBNA1
EBNA3A
EBNA3B
BZLF2
BTRF1
Cluster III (N= 8)
Mean Norm. Expression
Day
Genes:
BNLF1/LMP1
LMP2B
BALF5
LMP2A
BALF4
BALF3
BXLF2
BARF0

Figure S10

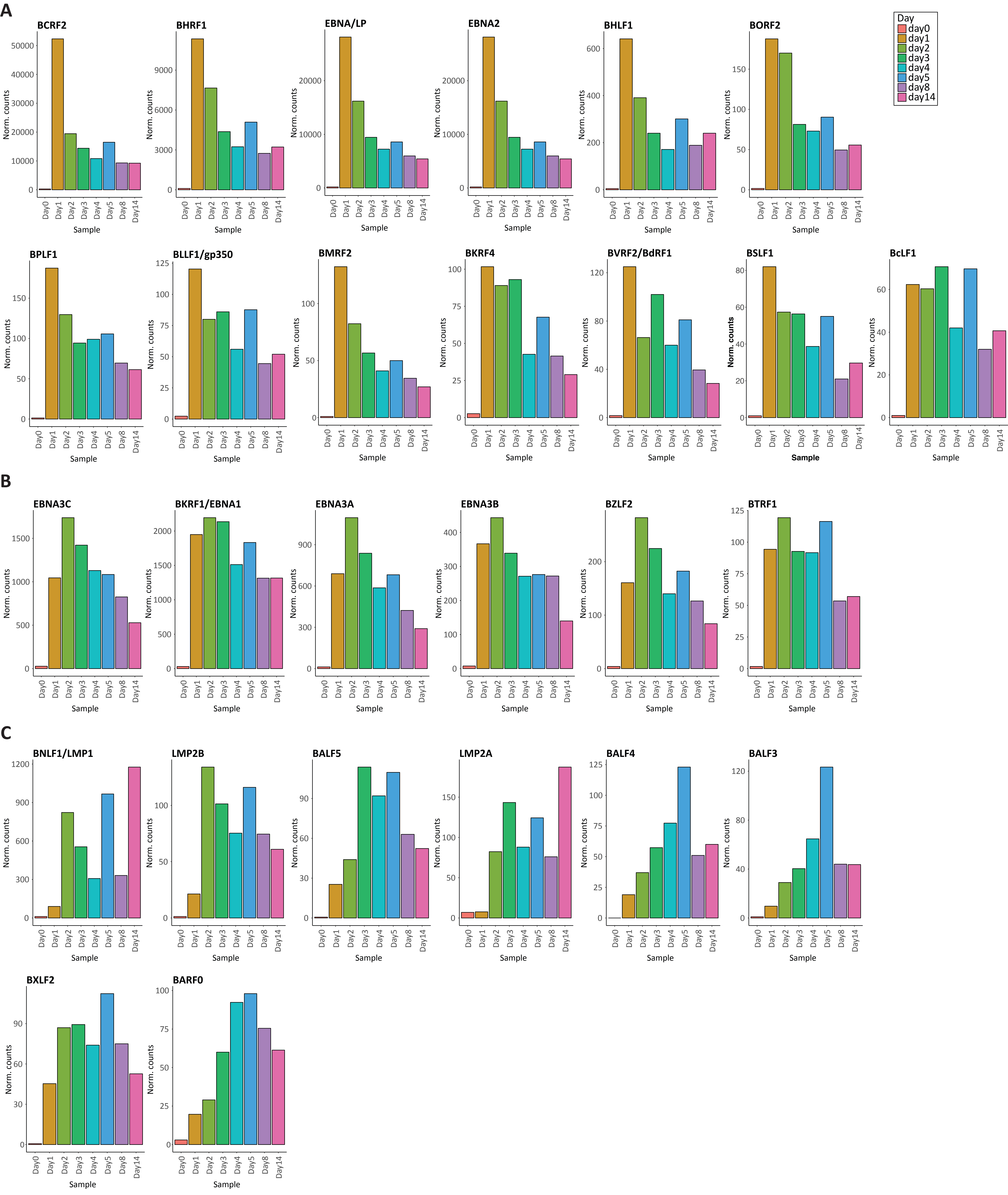
A
BCRF2
BHRF1
EBNA/LP
EBNA2
BHLF1
BORF2
Day
day0
day1
day2
day3
day4
day5
day8
day14
BPLF1
BLLF1/gp350
BMRF2
BKRF4
BVRF2/BdRF1
BSLF1
BcLF1
B
EBNA3C
BKRF1/EBNA1
EBNA3A
EBNA3B
BZLF2
BTRF1
C
BNLF1/LMP1
LMP2B
BALF5
LMP2A
BALF4
BALF3
BXLF2
BARF0
Norm. counts
Sample

Figure S11
